# Pangenome and Transcriptomic Analysis Revealed Distinct Virulence Genes Profiles and Infection Responses in *Shigella Sonnei* and *Shigella Flexneri*

**DOI:** 10.1101/2024.08.22.609085

**Authors:** Bao Chi Wong, Wei Yee Wee, Hock Siew Tan

## Abstract

Shigellosis, a major cause of diarrheal death, is predominantly caused by *Shigella flexneri* and *Shigella sonnei*. Notably, *S. sonnei* has begun to overtake *S. flexneri* in incidence worldwide. We conducted a comparative pangenome and transcriptomic analysis of these two species using the well-established model *Caenorhabditis elegans* as an infection model to explore this shift. Pangenome analysis of 46 *S. sonnei* and 111 *S. flexneri* strains revealed that *S. sonnei* has more core genes (44%) than *S. flexneri* (13.6%). Additionally, *S. sonnei,* but not *S. flexneri,* has core virulence genes. The transcriptomic study during *C. elegans* infection showed that *S. sonnei* significantly upregulates the ferric enterobactin outer membrane transporter (*fepA*) and other metabolic processes (sulfur, thiamine and arginine). In contrast, *S. flexneri* upregulates regulatory genes involved in biofilm formation, transcription, cellular processes, and cell communication. These findings suggest that *S. sonnei*’s ability to upregulate specific genes, including *fepA,* may confer a competitive advantage over *S. flexneri*. These findings provide a comprehensive understanding of the genetic and functional adaptations that may contribute to the global epidemiological shift from *S. flexneri* to *S. sonnei*, underscoring the need for further research into the mechanisms driving *S. sonnei*’s increased virulence and adaptability.

## Introduction

Shigellosis is an intestinal infection caused by *Shigella* spp., with symptoms such as watery or bloody diarrhea, and vomiting. It is the second leading cause of diarrheal death, and it is more prevalent in children under five years of age [1]. While this disease is usually self-limiting, it is highly infectious as 10 viable *Shigella* is sufficient to cause shigellosis [2]. It spreads through food or water contamination [3], or through direct fecal-oral contact [4]. Reports from recent decades revealed the increased prevalence of *Shigella sonnei,* which is slowly overtaking *Shigella flexneri* as the cause of shigellosis, especially in low- and middle-income countries (LMIC) [1, 5–8]. Multiple studies have hypothesized the reasons behind this shift. These hypotheses include the potential cross-immunization from *Plesiomonas shigelloides* [9], the ability of *S. sonnei* to outcompete *S. flexneri* [10], and the ability of *S. sonnei* ability to acquire antimicrobial genes from commensal and pathogenic bacteria [5].

*S. sonnei* is distinct from *S. flexneri* in terms of their pathogenesis. For instance, their O-antigens are different. *S. flexneri* has at least 19 serotypes, divided based on the antigenic determinants of the O-antigen of the lipopolysaccharide (LPS) [11]. In contrast, *S. sonnei* only has one serotype, which is distinct from other *Shigella* species. Instead, it has a higher similarity to the O-antigen of *P. shigelloides,* which may be the source of *S. sonnei* O-antigen [9, 12]. *S. sonnei* O-antigen, along with their group 4 capsules (G4C), plays a role in immune evasion by masking their epitopes from host antibodies [13]. Additionally, *S. sonnei,* which induces less pyroptosis and less cytosolic escape compared to *S. flexneri,* reduces its pathogenicity and inflammatory potential, leading to better spread across the host system [13]. *S. flexneri,* which is more aggressive, invades the macrophages via Type III Secretion System (T3SS) and delivers effector proteins that help bacterial proliferation within the host cell [14, 15].

*Shigella’s* pangenome regarding its phylogenetics and dissemination had been analyzed previously. In terms of dissemination, an analysis from 132 globally distributed *S. sonnei* isolates suggested that the current pandemic spread of *S. sonnei* can be linked to a single, multidrug-resistant lineage in the seventeenth century but has since segregated into at least four different lineages [16]. Other studies looking into *Shigella* focused on antimicrobial resistance [17, 18]. A few studies investigated the *Shigella’s* virulence genes and distribution among the different *Shigella* species [17, 19, 20]. These studies only analyzed common *Shigella* virulence factors, such as T3SS genes, aerobactin, enterobactin, and exotoxins that are within the Virulence Factor Database (VFDB) [21]. However, *S. sonnei* has been reported to have additional virulence genes not present or functional within *S. flexneri.* This includes a putative multivalent adhesion molecule (MAM) needed for attachment and invasion of host cells, but it is truncated within *S. flexneri* [22]. Additionally, the G4C operon (*ymcDCBA, yccZ, etp, etk*) that plays a role in improving the spread of *S. sonnei* are inactivated in *S. flexneri* due to a 14 bases deletion [13]. Lastly, *S. sonnei* has a functional Type VI Secretion System (T6SS) that provides a competitive advantage against other enteropathogens such as *S. flexneri* and *Escherichia coli* [10]*. S. flexneri*, however, had lost its T6SS due to missing genes, or gene inactivation. As these reported virulence factors are relatively novel and not common in all *Shigella* species, they are not included in the abovementioned databases. Thus, including these virulence factors in the pangenome analysis may help explain how *S. sonnei* is starting to overtake *S. flexneri* as the cause of the disease.

The differences between transcriptomic responses of *S. sonnei* and *S. flexneri* during host infection can also be a potential cause of the etiological shift. However, there are only a few studies looking into the response of *Shigella* during infection, and none had compared the responses between *S. sonnei* and *S. flexneri* during host infection. One study reported the transcriptomic response of *S. sonnei* during the zebrafish infection [23], while another reported the transcriptomic response of *S. flexneri* during adhesion to epithelial cells [24]. Thus, a comparative study of the transcriptome of *S. sonnei* and *S. flexneri* during host infection would be useful to observe any differences in bacterial responses.

In this study, we demonstrated that there are minimal differences between two representative strains, *S. sonnei* ATCC 29930 and *S. flexneri* ATCC 12022 during *in vitro* comparison. A pangenome analysis of *S. sonnei* and *S. flexneri* revealed that *S. sonnei* has more core genes than *S. flexneri.* The presence of common and newly reported virulence factors was identified in all strains using BLASTp, and some virulence genes such as G4C, MAM, T6SS and enterobactin synthesis and transport are closely related to *S. sonnei*. We also discovered that *S. sonnei* and *S. flexneri* employ different strategies during *Caenorhabditis elegans* infection using transcriptomics to compare the differences in gene expression between these two species.

## Methodology

### *In vitro* characterization of *Shigella sonnei* and *Shigella flexneri*

*Shigella sonnei* ATCC 29930 (*S. sonnei* 29930) and *Shigella flexneri* ATCC 12022 (*S. flexneri* 12022) were obtained from American Type Culture Collection (ATCC). Unless stated otherwise, all bacterial strains were grown on Luria-Bertani (LB) agar (Himedia, India) and incubated overnight at 37°C. Overnight cultures were grown in LB broth (Himedia, India) and incubated overnight at 37°C with 200 rpm shaking. The growth curves of both bacteria strains were determined by inoculating 50 ml of LB broth in a 250 ml conical flask with an aliquot of overnight bacterial culture. The conical flasks were incubated at 37°C with 200 rpm shaking, and the optical density (OD) 600nm of the culture was measured every 30 minutes until the bacterial culture reached plateau. Three technical and biological replications were done. The graph of OD 600nm against time was plotted.

### Pangenome analysis of 46 *S. sonnei* strains and 111 *S. flexneri* strains

The genome sequences of *Shigella sonnei* and *Shigella flexneri* were retrieved from public databases (accessed on 19 November 2023). A total of 46 *S. sonnei* and 111 *S. flexneri* genomes were included in this analysis (Supplementary Table 1). The chromosomes and plasmids of each strain were separately annotated with Prokka v1.14.5 [25]. The --protein parameter was used with *S. sonnei* Ss046 and *S. flexneri* 2a strain 301 GenBank files. Pangenome analysis was performed using the PGAP software [26]. The protein sequences of all strains (46 *S. sonnei* and 111 *S. flexneri*) were used as the input. Orthologs among these strains were searched using the Gene Family (GF) method. The protein sequences of each strain were mixed and marked with the strain identifiers. BLASTALL was first performed among the protein sequences with the minimum score value and E-value in BLAST as 50 and 1e-8 respectively. The filtered BLAST results were clustered by MCL algorithm. To group the same genes into the same cluster, the global match region must have at least 50% of the longer gene protein sequence and 95% sequence identity. Among the bacterial species, the genes present in all strains are considered core genomes, while the genes only present in one strain are considered unique genomes. The openness of the pangenome was determined using Heaps’ law [27], formulated as n = κN^γ^, where n is the pangenome size, N is the number of genomes, and κ and γ are the fitting parameters. If the exponent γ is less than 0, the pangenome is closed, while if γ is more than 0, the pangenome is open.

### Identification of virulence genes

The amino acid sequences of the 118 virulence genes were compiled by combining the *S. sonnei* and *S. flexneri-*specific virulence factors from VFDB, and other virulence factors that have reported differences between *S. sonnei* and *S. flexneri* (Supplementary Table 2). The annotated protein sequences of *S. sonnei* and *S. flexneri* were searched against the compiled virulence genes through BLASTp. The BLAST hits were further filtered with percent identity (piden) of more than 90%; query sequence coverage (qcov) of more than 90%, and the e-value of less than 1e-5. All heatmaps were constructed using R version 4.2.2 (http://www.r-project.org), with ggplot2 [28].

### Nematode killing assay and bacterial colonization assay of *Caenorhabditis elegans*

The *C. elegans* wild-type N2 Bristol strain was cultured on nematode growth medium (NGM) agar plates seeded with UV-killed *E. coli* OP50 and maintained at room temperature (22°C) [29]. *E. coli* OP50 is the standardized food source for *C. elegans* in laboratory conditions. *C. elegans* killing assay was performed as described previously [30, 31] with slight modifications. 100 µl of *S. sonnei* 29930 or *S. flexneri* 12022 overnight cultures were seeded on 60 mm diameter NGM agar plate and incubated at 37°C overnight. Plates were cooled to room temperature before use. The worms at L4 stage grown on UV-killed *E. coli* OP50 were transferred to the seeded NGM plates and incubated at 25°C. Worms were transferred to another seeded plate every two days for the first five days, and every three days henceforth to separate the adults from their progeny. *E. coli* OP50 was used as a negative control. Approximately 50 worms were used in the killing assay per replicate, and three biological replicates were done for statistical analysis. The number of live and dead worms was scored every 24 hours for 12 days. The worms were considered dead if they did not respond when touched by the worm pick. Worms that crawled up the walls of the plate or burrowed into the agar were censored.

To examine the bacterial accumulation within the nematodes, the bacterial colonization assay was performed as described previously [31]. The plasmid pUltra-RFP [32] containing kanamycin resistance was transformed into *S. sonnei* and *S. flexneri* through heat-shock/calcium chloride procedure [33]. The *C. elegans* were exposed to the bacteria similarly as above. At every 24 hours, 10 worms were transferred to 5 µl of M9 buffer containing 25 mM levamisole and 1mg/ml of gentamycin (MLG) on an NGM agar plate to remove the surface bacteria. After the MLG dried, the worms were transferred to a new 5 µl of MLG, and this washing step was performed 3 more times. The worms were transferred to 50 µl of sterile PBS + 1% Triton X-100 and mechanically disrupted using a sterile micropestle. The worm lysates were diluted and plated on nutrient agar containing 50 µg/ml kanamycin. The plates were incubated overnight at 37°C, and the colonies were quantified and used to calculate the number of bacteria per nematode.

### *C. elegans* infection and RNA extraction

RNA extraction of the *C. elegans* – *Shigella* was performed as previously described [34]. At 10 minutes (0 hours post infection, hpi) and 72 hpi, 100 worms were collected in 500 µl of M9 + 25 mM levamisole (ML). The worms were washed twice with ML and incubated in MLG for 1 hour at room temperature. After an additional wash in ML, 1 ml of Tri-RNA (Favorgen) was added, and the worms were flash-frozen in liquid nitrogen and stored at −80°C until RNA extraction. Briefly, the worm suspensions were freeze-thawed multiple times to break the worm cuticle. After adding 200 µl of chloroform (inverted and left to sit at room temperature for 5 minutes), the samples were centrifuged at 13000 rpm for 15 minutes at 4 °C. The RNA-containing upper phase was transferred to a new microcentrifuge tube. Total RNA extraction was done using the Monarch Total RNA Miniprep kit (New England Biolabs, USA) according to the instructions provided by the manufacturer, and on-column DNase treatment was performed.

### RNA sequencing and transcriptome analysis

All RNA samples were sent to AGTC Genomics (Malaysia) for sequencing. The samples were processed using Illumina Stranded Total RNA Prep and Ribo-Zero Plus kit (Illumina, CA, USA) according to the manufacturer’s instructions. The samples were then sequenced using Illumina NovaSeq 6000 platform, with the conformation of 150 bp paired ends. The reads were trimmed using Trimmomatic v0.39 [35] and first mapped to the WBcel235 *C. elegans* genome using HISAT2 v2.2.1 [36]. The unmapped reads were then mapped to their respective bacterial genome using HISAT2, and read counts were quantified using HTseq v0.11.2 [37]. The summary statistics of the transcriptomic analysis can be found in Supplementary Table 3. Data normalization and differential abundance analyses were conducted using DESeq2 [38] with a cutoff of false discovery rate (FDR) of < 0.05, and fold change (FC) of > 2. Heatmaps and charts were generated using R version 4.2.2 with ggplot2 [28]. The common genes between the two species were obtained using ProteinOrtho [39]. Only genes with one ortholog in both species were analyzed. Gene Ontology functional pathways enrichment was performed using ShinyGo v0.80 [40]. The Clusters of Orthologous Groups (COGs) of the genes were determined using EggNOG v5.0 [41].

### Statistical analysis

To statistically test the difference in virulence genes in *S. sonnei* and *S. flexneri,* the Mann-Whitney U test was performed. The nematode survival was analyzed using the Kaplan-Maier method, while the significant differences between survival curves were determined in a log-rank (Mantel-Cox) test using GraphPad Prism, version 10 (GraphPad Software, San Diego, CA, USA). The bacterial colonization of different bacteria was analyzed using a two-way analysis of variance (ANOVA) followed by Sidak multiple comparison tests across the same time point (ns = non-significant). Unless stated otherwise, *p* values < 0.05 are considered statistically significant.

## Results and Discussion

### Strain comparison between *S. sonnei* and *S. flexneri* revealed similar growth but distinct genome profile

While *S. flexneri* remains the leading cause of shigellosis, *S. sonnei* has begun to overtake *S. flexneri* as the etiological cause of the disease [5, 17]. To study the differences between these two species, *S. sonnei* 29930 and *S. flexneri* 12022 were selected as the representative strains for analysis. These two strains were chosen as they were obtained from ATCC, commonly used for infectious disease research and readily available in our lab. *S. sonnei* 29930 is also a part of the Walter Reed Army Institute of Research (WRAIR) collection of virulent strains.

*In vitro* growth curve study demonstrated that these two strains have similar growth under laboratory conditions (Figure 1a). Next, the bacteria whole genome sequences were obtained from the public databases. To compare the genes and coding sequences (CDS) in their genome, the common genes between these two species were obtained using ProteinOrtho. This tool identifies orthologous and co-orthologous proteins using a BLAST-based approach. *S. sonnei* 29930 and *S. flexneri* 12022 have 5364 and 5269 total genes respectively (Figure 1b), where 3366 genes have a single ortholog in each species. There are 256 and 345 genes with multiple orthologs in *S. sonnei* 29930 and *S. flexneri* 12022 respectively. Both strains have similar number of strain-specific genes, with 1742 and 1558 genes specific to *S. sonnei* 29930 and *S. flexneri* 12022. The genome sequencing summary statistics were also compared (Figure 1c). While *S. sonnei* 29930 has a complete genome with a single plasmid, *S. flexneri* 12022 has a circularized chromosome, 4 circularized plasmids, and multiple linear contigs. An important virulence plasmid, pINV (>200kb plasmid) is important for *Shigella* virulence as it encodes genes associated with the ability to invade epithelial cells in the gastrointestinal tract [42, 43]. However, these two strains do not have this plasmid in their genome.

**Figure 1:**
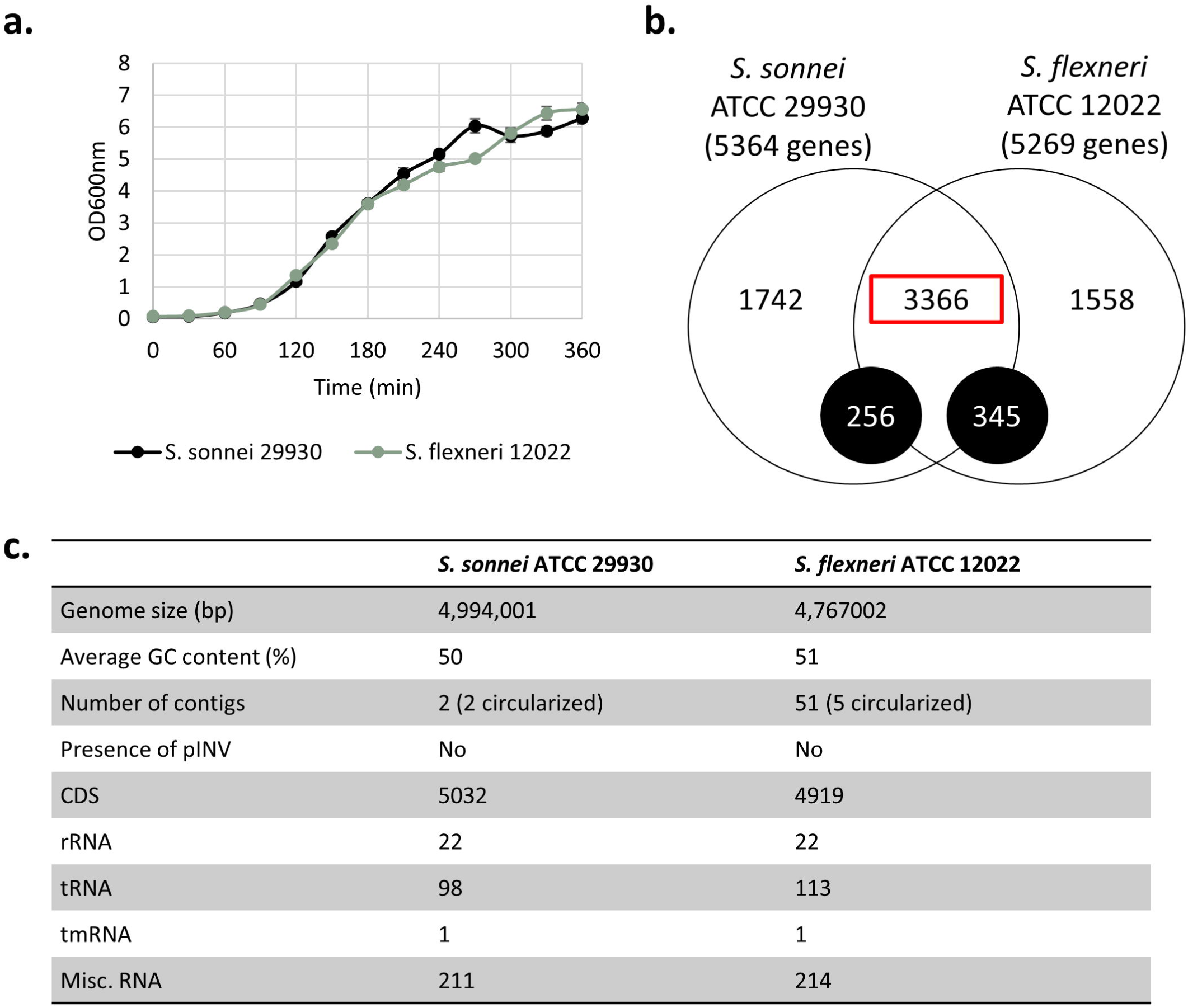
Comparison between *S. sonnei* 29930 and *S. flexneri* 12022. **a.** *In vitro* growth curve of the two strains. **b.** Venn diagram of the common genes between the two strains. The black circle denotes the common genes with multiple orthologs, while the red square denotes the common genes with a single ortholog in both strains. **c.** Summary statistics of the genomes of the two strains.

### *S. sonnei* has more core genes than *S. flexneri*

A pangenome analysis using 46 *S. sonnei* and 111 *S. flexneri* genomes from public databases was performed to elucidate the species differences between *S. sonnei* and *S. flexneri*. The pangenome analysis of both species was performed on the chromosome and plasmid levels using PGAP. Firstly, the status of the pangenomes was determined to be open or closed using Heaps’ Law [27]. An open pangenome indicates that new genes would be found as the number of strains increases, while a closed pangenome indicates that no more new genes would be found after a certain number of genomes. Both *S. sonnei* and *S. flexneri* have an open pangenome, with the γ values of 0.644 and 0.585, respectively, using the Heaps’ Law (Figure 2a and 2b). This is similarly reported in other studies on *Shigella* [18, 44] and *E. coli*, a close relative of *Shigella* [45].

**Figure 2:**
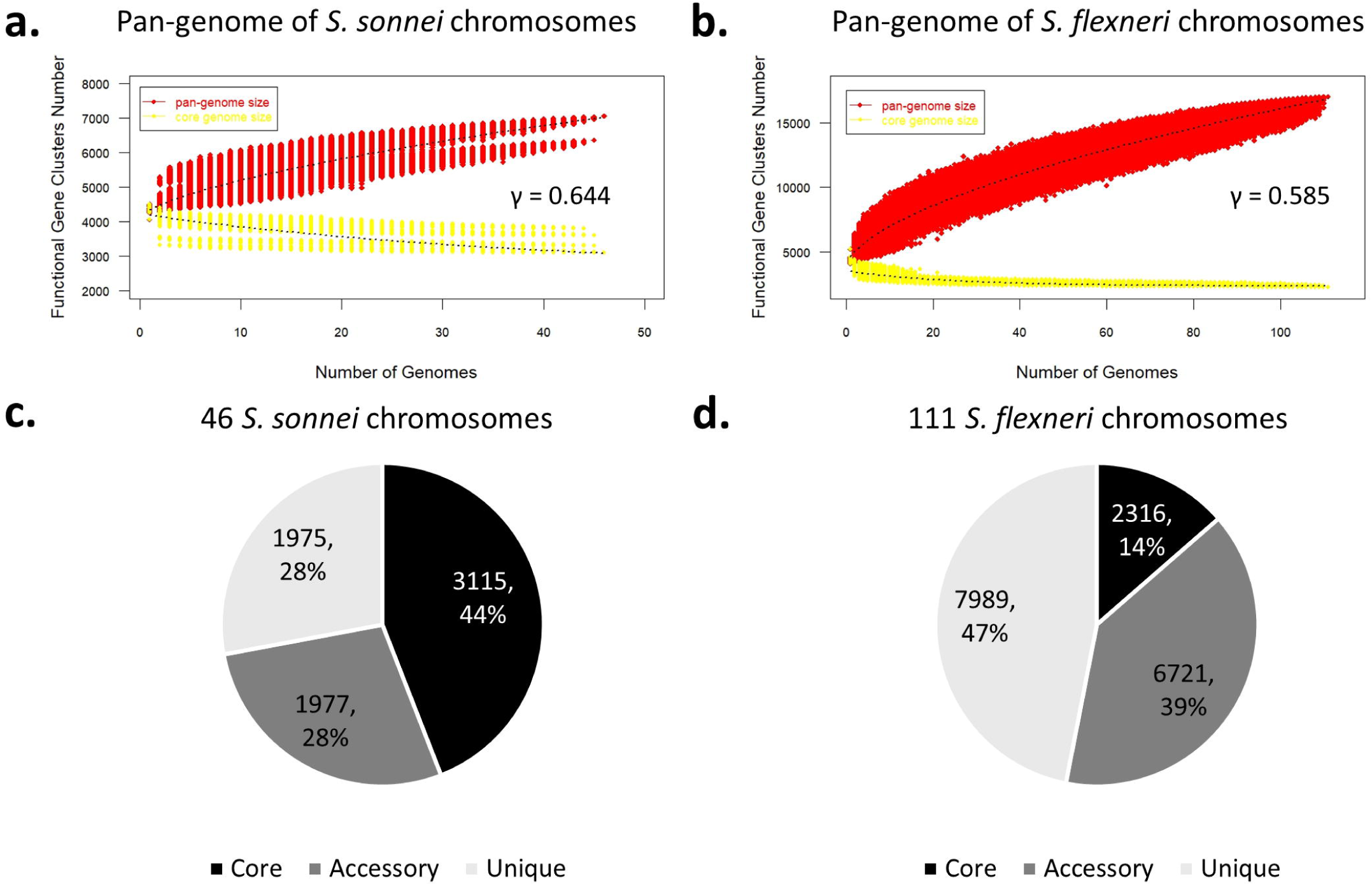
Pangenome analysis of the chromosomes of all 46 *S. sonnei* and 111 *S. flexneri* strains. **a.** Gene accumulation curves of the pangenome (red) and core genome (yellow) of 46 chromosomes of *S. sonnei,* and **b.** One hundred eleven chromosomes of *S. flexneri*. The boxes show the γ values, showing that both pangenomes are open. **c.** Pie chart depicting the number of core, accessory and unique genes of *S. sonnei* and **d.** *S. flexneri*.

All gene clusters were categorized into 3 groups: core (present in all the strains), unique (present in only 1 strain), and accessory (present in >2 strains, but not all strains) (Figure 2c and 2d). In the 46 *S. sonnei* chromosomes, a total of 7067 gene clusters were identified, where 44% (3115 genes) are core, 27.95% (1977 genes) are accessory, and 27.98% (1975 genes) are unique. In the 111 *S. flexneri* chromosomes, there are 17026 total gene clusters, where 13.6% (2316 genes) are core, 39.47% (6721 genes) are accessory, and 46.92% (7989 genes) are unique. *S. sonnei* has more core genes within its pangenome (44% of *S. sonnei* genes compared to 13.6% of *S. flexneri* genes). In comparison, *S. flexneri* has more unique genes (47% of *S. flexneri* genes compared to 28% of *S. sonnei* genes). However, another pangenome study on 82 *S. sonnei* strains and 106 *S. flexneri* strains revealed that 6% of *S. sonnei* genes are core genome, while 15% of *S. flexneri* genes are core genome [17]. This may be due to the different number of strains used, where twice the number of *S. sonnei* strains were analyzed. Other studies on *S. flexneri* pangenome reported a higher percentage of core genes with a lower number of *S. flexneri* strains. For example, 66% of genes were identified as core across 15 *S. flexneri* genomes [19], while another analysis on 11 *S. flexneri* strains identified 60% core genes [44]. Nevertheless, our study demonstrated that generally, the distribution of core, accessory, and unique gene clusters differed between these two species. Our study is also the first to perform a pangenome analysis on all the currently available complete genomes of *S. sonnei* and *S. flexneri* on public databases.

### The prevalence of pINV and the plasmid-encoded virulence genes are similar between *S. sonnei* and *S. flexneri*

This study investigated the prevalence of virulence genes in *S. sonnei* and *S. flexneri* in each strain. The protein sequences of 118 virulence genes were curated from VFDB and other studies of *S. sonnei* virulence, and their presences in the genomes are determined using BLASTp. These genes are considered present and functional when there is >90% identity and >90% coverage. The virulence genes in this study are broadly categorized into plasmid-encoded virulence genes (71 genes) and chromosomal virulence genes (47 genes).

The plasmid-encoded genes are categorized into colicins, T3SS, exoenzymes, and others. Except for the 12 colicin genes, these virulence genes are found on the large pINV, as previously described [14]. It is reported that the loss of this plasmid in *Shigella* is common during laboratory culturing of the *Shigella* strains before genomic DNA extraction. In particular, *S. sonnei* has a higher chance of losing this plasmid due to the loss of two toxin-antitoxin (TA) systems and a single amino acid polymorphism in another TA system [46]. In this study, the presence of pINV is indicated by a >200kb plasmid. Around 30% of both species possess a 200kb plasmid (Figure 3a; detailed heatmap of plasmid-encoded virulence genes in Supplementary Figure 1 for *S. sonnei,* and Supplementary Figure 2 for *S. flexneri*). The remaining strains either do not have plasmids at all, or only have smaller size plasmids. Around 95% of strains with pINV have the exoenzymes (*icsP/sopA, sepA*), *icsA/virG, virF, virK,* and the T3SS genes (Figure 3b). In contrast, less than 20% of strains without pINV have these genes. These pINV-encoded virulence genes are not associated with the *Shigella* species, but rather the presence of this large virulence plasmid. Interestingly, among the T3SS genes, *ipaJ* has a low prevalence within *S. sonnei* and *S. flexneri.* BLASTp revealed that the *ipaJ* gene was truncated in 7 *S. sonnei* and 38 *S. flexneri* strains, where 32-58 amino acids were lost. IpaJ is one of the T3SS effectors, and its role as a cysteine protease alters the protein trafficking in host cells, thereby preventing autophagy [47]. Expression of *ipaJ* is also linked to the increased *S. flexneri* bacterial load within a host cell [48]. No studies have been performed to investigate the prevalence of *ipaJ* in *Shigella,* and there were no previous reports of truncation in this protein.

**Figure 3:**
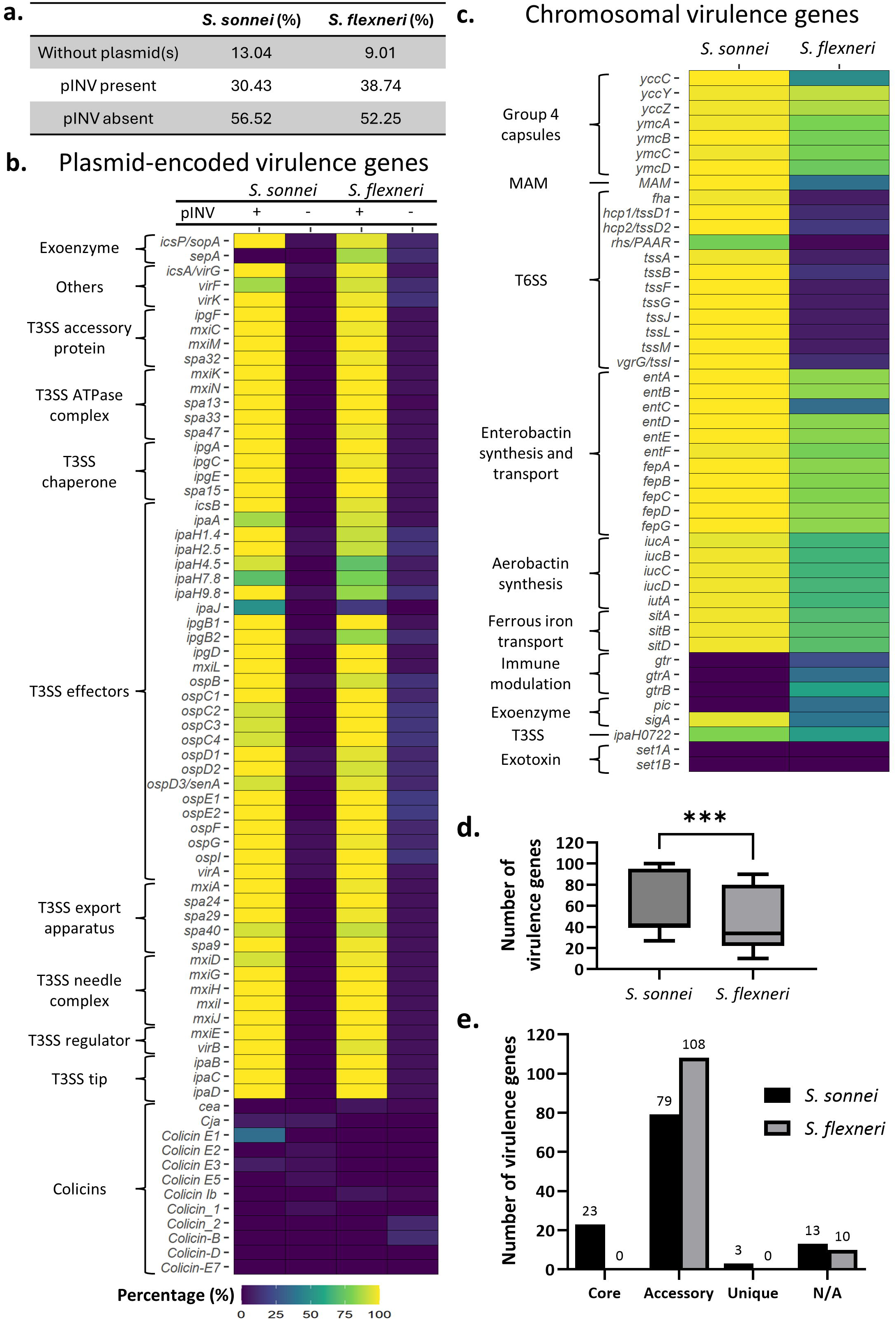
Prevalence of pINV and virulence genes in *S. sonnei* and *S. flexneri* identified using BLASTp. **a.** Prevalence of pINV or other plasmids in all *S. sonnei* and *S. flexneri* in this study. **b.** Prevalence of pINV-related virulence genes in the *S. sonnei* and *S. flexneri* strains. W indicates strains with pINV, while Wo indicates strains without pINV. **c.** Prevalence of chromosomal virulence genes in the *S. sonnei* and *S. flexneri* strains. All heatmaps are scaled according to the percentage (%) of strains with the virulence genes. (T3SS = Type III secretion system, MAM = multivalent adhesion molecule, T6SS = Type VI secretion system). **d.** Boxplot of the number of virulence genes in *S. sonnei* and *S. flexneri* (Mann-Whitney-U test was conducted, p-value *** indicates <0.0004). **e.** Number of core, accessory, and unique virulence genes in *S. sonnei* and *S. flexneri.* N/A are virulence genes not present in any *S. sonnei* or *S. flexneri* strains.

Lastly, colicins are rarely found in the plasmids of both *Shigella,* as they are present in less than 13% of the strains analyzed (Figure 3b). *S. sonnei*, but not *S. flexneri,* has been reported to use colicins to compete against other pathogens during gastrointestinal infection [49–51]. However, our analysis shows no apparent difference in the colicin’s prevalence between these two species.

### Previously reported *S. sonnei*-associated virulence genes (G4C, MAM, T6SS) are conserved in most *S. sonnei* strains, but not *S. flexneri*

There are 47 chromosomal virulence genes within *S. sonnei* and *S. flexneri* (Figure 3c; detailed heatmap of chromosomal virulence genes in Supplementary Figure 3 for *S. sonnei,* and Supplementary Figure 4 for *S. flexneri*). These genes are categorized into the group 4 capsules (G4C), multivalent adhesion molecule (MAM), type VI secretion system (T6SS), enterobactin synthesis and transport genes, aerobactin synthesis genes, ferrous iron transport genes, immune modulation, exoenzymes, exotoxins, and one gene from T3SS (*ipaH0722*). 62% of these genes (23/37) are core genes in *S. sonnei*, but none are in *S. flexneri.* Among the 10 categories of chromosomal virulence genes, 6 categories are present in more *S. sonnei* than *S. flexneri*.

The genes in the G4C operon (*yccC, yccY, yccZ, ymcA, ymcB, ymcC,* and *ymcD*), the MAM gene, and the T6SS (*fha, hcp1, hcp2, rhs, tssA, tssB, tssF, tssG, tssJ, tssL, tssM,* and *tssI*) have been associated with *S. sonnei,* but not *S. flexneri* virulence [10, 13, 22]. Indeed, BLASTp results revealed that there are more *S. sonnei* strains with G4C, MAM and the T6SS genes than *S. flexneri* (Figure 3c). More than 97% of *S. sonnei* possess the complete G4C operon, 100% have the MAM gene, and more than 78% have the T6SS genes in this study. In contrast, only 48% of *S. flexneri* possess the complete G4C operon, 37% have the MAM gene, and less than 16% have the T6SS genes. The loss of these genes in the *S. flexneri* genome may be due to deletion or mutations in the gene that renders the gene(s) non-functional. All genes in the G4C operon must be functional for the successful formation of the capsule [52]. However, 44 strains of *S. flexneri* have the 14 bases gap in their *yccC* DNA sequence, while 12 strains have a truncated *yccC*. This deletion can lead to a frame-shift mutation, leading to the inactivation of *yccC* and thus the inactivation of the G4C operon [13]. The MAM protein is needed for attachment and invasion of host cells by *S. sonnei*, but it is heavily truncated in *S. flexneri* [22]. Indeed, BLASTp results revealed that among the *S. flexneri* strains without MAM, most only have 511 out of 877 amino acid sequences of the MAM protein. Lastly, the loss of the T6SS genes in the *S. flexneri* genome is quite prevalent due to gene deletion or inactivated by insertion sequences or stop codons [10]. The high prevalence of these genes in *S. sonnei* and not in *S. flexneri* indicates their importance in *S. sonnei* pathogenesis and highlights the species-specific differences between these two species.

### Enterobactin synthesis and transport, aerobactin synthesis, and ferrous iron transport genes are present in most *S. sonnei* and not in *S. flexneri*

Unlike the G4C, MAM, and T6SS, the enterobactin synthesis and transport (*ent* and *fep* genes), aerobactin (*iuc* genes and *iutA*), and ferrous iron transport genes (*sit* genes) have not been directly linked to *S. sonnei-*specific virulence. More than 95% of *S. sonnei* in this study have these virulence genes, but not *S. flexneri* (Figure 3c). BLASTp results revealed that around 20% of *S. flexneri* have a high number of mismatches in the Ent (EntA, EntB, EntE, EntF) and Fep (FepA, FepB, FepC, FepD, FepG) animo acid sequences. They also have truncated EntC, while EntD is not detected in these strains. Around 35% of *S. flexneri* strains do not have the aerobactin synthesis genes, while around 30% have a high number of mismatches in the Sit amino acid sequences. These three categories of genes are involved in iron uptake and utilization. Iron is a co-factor for essential enzymes involved in multiple biological processes such as gene regulation, central metabolism, and DNA replication [53]. Due to its importance, limiting the iron levels within the host is one of the innate immune mechanisms against bacterial pathogen [54]. To counter this response, pathogens including *Shigella* produce siderophores such as enterobactin and aerobactin to acquire iron. The siderophores are first synthesized by *ent* genes (enterobactin) and *iuc* genes (aerobactin) and transported out of the cell [55]. After scavenging ferric iron from the host, the ferric siderophores are taken up via FepA or IutA into the cell for iron utilization. By sequestering iron from the host, these siderophores promote bacterial survival during infection [56]. The Sit system is a part of the ferrous iron utilization system involved with iron uptake. One study reported that *S. flexneri* can depend solely on the Sit system for iron-uptake and form plaques on Henle cell monolayers [57]. The usage of different types of siderophores have been reported in *S. flexneri* and *S. sonnei.* For instance, *S. flexneri* does not utilize enterobactin as some genes were inactivated in *S. flexneri* due to frameshift mutations, insertion sequences or stop codons [58]. *S. sonnei* contains the genes for both aerobactin and enterobactin systems, but while it cannot synthesize its own aerobactin, they are able to utilize exogenous aerobactin [59]. These iron uptake and utilization genes are highly prevalent in *S. sonnei,* suggesting that *S. sonnei* may have conserved their iron uptake and utilization repertoire in their core genomes.

### The *Shigella* enterotoxin 1 genes are absent in both species, while *gtr* genes and exoenzymes are absent in either *S. sonnei* or *S. flexneri*

Multiple virulence genes are completely absent from *S. sonnei* or *S. flexneri,* or both. The *Shigella* enterotoxin 1 (ShET1) genes *set1a* and *set1b* are absent in all strains in this study. This differs from other reports which demonstrated that these two genes were detected through PCR in less than 10% of *S. sonnei* [60], and around 48-78% of *S. flexneri* [61, 62]. ShET1 is reported to be exclusively found in *S. flexneri* 2a [62, 63], explaining the genes’ absence in *S. sonnei* and other *S. flexneri* strains. However, their absence in the *S. flexneri* 2a strains analyzed here may be due to the discrepancy between PCR detection of the genes and BLASTp detection of the genome annotations.

Other virulence genes have a different prevalence profile between *S. sonnei* and *S. flexneri.* The glycosyltransferase genes (*gtr, gtrA* and *gtrB*) are present in the chromosomes of *S. flexneri* but not *S. sonnei.* These genes are involved in lipopolysaccharide (LPS) immune modulation and O-antigen modification [64]. They are used in serotype conversion for host immune evasion, a characteristic all *Shigella* share except *S. sonnei,* as they only have one serotype [12]. Other than that, both *S. sonnei* and *S. flexneri* have different prevalences of the exoenzymes genes, *Shigella* extracellular protein (*sepA*) in the pINV, and the protease involved in intestinal colonization (*pic*) and *Shigella* IgA-like protease homologue (*sigA*) in the chromosome. These 3 genes are also classified as SPATEs (serine protease autotransporters of Enterobacteriaceae) genes. *Shigella* utilize these genes to induce inflammation and damage the intestinal cells [65]. 95% of *S. sonnei* and 39% of *S. flexneri* have the *sigA* gene. The *sigA* gene is present in more *S. sonnei* strains (95%) than *S. flexneri* (39%). The *pic* and *sepA* genes are present only in *S. flexneri* strains (*pic:* 36%; *sepA:* 86%), as no *S. sonnei* strains have these genes in their genome (Figures 3b and 3c). The prevalence of *sigA, pic,* and *sepA* are similarly reported in other studies [60, 61, 66, 67]. SigA has enterotoxin activities similar to ShET1; Pic promotes better access to the epithelium cell through mucin cleavage, while SepA induces fluid accumulation [14]. As these exoenzymes play different roles during infection, their distinct prevalence in *S. sonnei* and *S. flexneri* suggest that their role has different importance in these two species during pathogenesis.

### *S. flexneri*, but not *S. sonnei,* has *gtrA, gtrB,* ferrous iron transport genes, aerobactin genes, and *virK* encoded in both chromosome and plasmids

*S. flexneri*, but not *S. sonnei,* has virulence genes encoded in both chromosome and plasmids (Supplementary Figure 5). For instance, *S. flexneri* 1a strain 0228 encodes *gtrA* in its plasmid instead of the chromosome. It also has a duplicated *gtrB* in its chromosome and its plasmids. The aerobactin genes (*iucA, iucB, iucC,* and *iucD*) are encoded in the chromosomes of *S. flexneri* and *S. sonnei,* but some *S. flexneri* strains encode these genes in their plasmids. Other than that, 3 *S. flexneri* strains possess two copies of the ferrous iron transport genes (*sitA, sitB,* sitD) in their genome - one in their chromosome and one in their plasmids. Other *S. flexneri* strains encode the *sit* genes in their plasmids instead of their chromosomes. VirK, a cytoplasmic polypeptide associated with *Shigella* virulence, is commonly located in the pINV. However, 2 *S. flexneri* strains encode this gene in their chromosomes, while 20 *S. flexneri* strains have two copies of *virK* in their genome (one in their chromosome and one in their plasmids). It is unclear why the *iut*, *sit,* and *virK* genes in some *S. flexneri* are encoded in both the chromosome and plasmid. However, an enteroaggregative *E. coli* has *virK* in both its chromosome and plasmid, indicating that this is not limited to *S. flexneri* [68].

### *S. sonnei* has more virulence genes than *S. flexneri,* and more core virulence genes than *S. flexneri*

Among the 118 virulence genes analyzed in this study, *S. sonnei* has significantly more virulence genes than *S. flexneri* (Figure 3d, p-value <0.0004), with 57 genes and 47 genes, respectively*. S. sonnei* has 19% (23/118 genes) of these virulence genes in their core genome, but *S. flexneri* do not have any core virulence genes (Figure 3e). Instead, 91.53% (108/118 genes) of the virulence genes in *S. flexneri* are accessory genes. This suggests that these virulence genes are more conserved in *S. sonnei* as they are encoded in almost all strains, whereas the virulence genes in *S. flexneri* are more diverse. To our knowledge, this is the first study that compared the amount of virulence genes between *S. sonnei* and *S. flexneri,* which demonstrated that *S. sonnei* possess more virulence genes in their genome than *S. flexneri*.

### *S. sonnei* is more virulent than *S. flexneri* towards *Caenorhabditis elegans*

To determine if the higher number of virulence genes translates to higher *in vivo* virulence, the nematode *C. elegans* was used as an animal model for *Shigella* infection using *S. sonnei* 29930 and *S. flexneri* 12022 strains, a representative strain for *S. sonnei* and *S. flexneri* respectively. Firstly, the presence of the virulence genes within these two strains was determined (Figure 4a and 4b). They share 24 common virulence genes, while 67 genes are absent in both strains (Figure 4a). There are 15 virulence genes (including the MAM and T6SS genes) only found in *S. sonnei* 29930, while 10 genes (*gtr* genes and some T3SS genes) are only found in *S. flexneri* 12022. Generally, both strains have aerobactin, exoenzymes, enterobactin, ferrous iron transport, and G4C genes (Figure 4b). As mentioned previously (Figure 1c), these two strains do not have the pINV, and thus do not have any pINV-encoded virulence genes.

**Figure 4:**
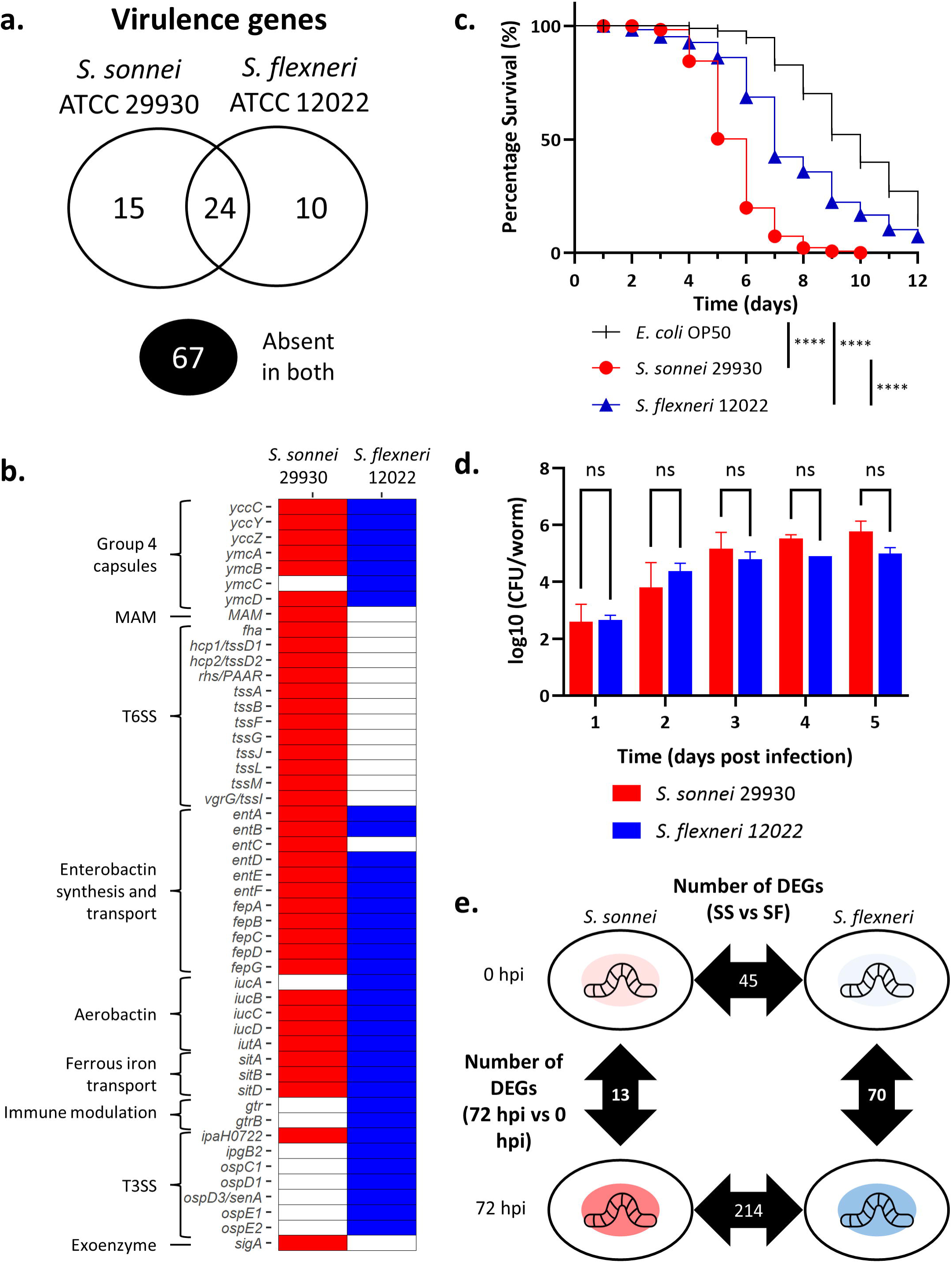
*C. elegans* infection by *S. sonnei* 29930 and *S. flexneri* 12022. **a.** Venn diagram of the virulence genes presence in *S. sonnei* 29930 and *S. flexneri* 12022. Black circle denotes the virulence genes that are not present in either of these two strains. **b.** Presence of virulence genes in *S. sonnei* 29930 (red) and *S. flexneri* 12022 (blue). **c.** Killing of *C. elegans* infected by *S. sonnei* 29930 (red circle), *S. flexneri* 12022 (blue triangle), and *E. coli* OP50 (black tick, negative control). The non-parametric log-rank (Mantel-Cox) test was used to analyze the survival curve (p-value **** p<0.0001). Killing assay data combines three independent experiments with around 60 worms each. d. Bacterial colonization assay of *C. elegans* infected by *S. sonnei* 29930 (red) and *S. flexneri* 12022 (blue). Bacterial colonization data combines three independent experiments with 10 worms each. Error bars represent the standard error of the mean (SEM). Statistical significance was analyzed using a two-way analysis of variance (ANOVA) followed by Sidak multiple comparison tests across the same time point (ns = non-significant). **e.** Schematic diagram of the infection experimental design and the identified differentially expressed genes (DEGs) between each comparison. The RNA transcripts of the *Shigella-C. elegans* infection were obtained from *S. sonnei* and *S. flexneri* infection at 0 hpi and 72 hpi.

Both strains are virulent towards the nematodes, as the survival curves are significantly different from the negative control (Figure 4c). The two strains are also significantly different from each other. *S. sonnei* 29930 needed five days to kill 50% of the nematode population, while *S. flexneri* 12022 needed seven days, indicating that *S. sonnei* 29930 is comparatively more virulent than *S. flexneri* 12022. This is similarly reported in a prior study on zebrafish conducted at 28.5 °C (the standard zebrafish maintenance temperature) [23]. However, both strains take a longer time compared to other reported *Shigella* strains in reducing the nematode population to 50%. Studies on *Shigella-C. elegans* infection reported that virulent *S. flexneri* and *S. sonnei* kill 50% of the worm population within 1 or 2 days [69, 70]. This may be due to the lack of pINV in this study. Nevertheless, these two strains of *Shigella* are virulent towards *C. elegans* at 25 °C.

Since the difference in virulence may be due to different bacterial loads within the nematode during infection, a bacterial colonization assay was performed to determine the bacteria colony forming unit (CFU) within *C. elegans* (Figure 4d). No significant difference in the CFU/nematode was observed between the two *Shigella* during infection at all 5-time points, suggesting that the differences in virulence observed are most likely due to their genomic and transcriptomic differences and independent of the differences in bacterial load.

### *S. sonnei* and *S. flexneri* did not upregulate their virulence genes during *C. elegans* infection

*S. sonnei* 29930 has a slightly higher number of virulence genes than *S. flexneri* 12022 (Figure 4a). In addition, it is more virulent than *S. flexneri* 12022 towards *C. elegans* with similar bacterial load within the nematode’s guts. Thus, the role of these virulence genes during infection were investigated through transcriptomics analysis to determine if their gene expressions are induced during infection. The nematodes were exposed to *S. sonnei* 29930 or *S. flexneri* 12022 for 72 hours, and the total RNA was extracted from the worms. This time point was chosen as the nematodes contain the maximum bacterial load while causing minimal death in the nematodes, allowing the analysis of early *Shigella* pathogenesis. As a control, the nematodes exposed to the bacteria for 10 minutes were also harvested (0 hpi) before the bacteria established an infection. Significantly differentially expressed genes (DEGs) were identified using the threshold log2(fold change) > |1| and adjusted p-value of < 0.05. The principal component analysis (PCA) was performed on the *S. sonnei* and *S. flexneri* data, confirming that the biological replicates are similar and cluster together (Supplementary Figure 6). Multiple differential gene expression analyses were performed between the 2-time points of infection (72 hpi vs 0 hpi) in *S. sonnei* 29930 and *S. flexneri* 12022 separately and between the two species at the same time of infection (*S. sonnei* vs *S. flexneri*) (Figure 4e).

There are 13 DEGs in *S. sonnei* 29930 at 72 hpi vs 0 hpi (Figure 5a; Supplementary Table 4). In contrast, 70 DEGs are identified in *S. flexneri* 12022 at similar conditions (Figure 5b; Supplementary Table 5). There are no virulence genes among the 13 genes differentially regulated by *S. sonnei*. In contrast, *S. flexneri* downregulates two aerobactin genes *iucA* and *iucD* at 72 hpi (Figure 5b, highlighted in red boxes). The *iucABCD* operon encodes for the enzymes required for aerobactin synthesis, which facilitates *S. flexneri* extracellular growth, but not intracellular growth [71]. However, *S. flexneri* is more suited to an intracellular lifestyle than *S. sonnei* as they are better at epithelial cells and macrophage invasion [72], which may explain why these aerobactin genes are downregulated in *S. flexneri.* This study revealed that although the virulence genes may be present in the *Shigella’s* genome, they are not necessarily highly expressed or differentially expressed during infection.

**Figure 5:**
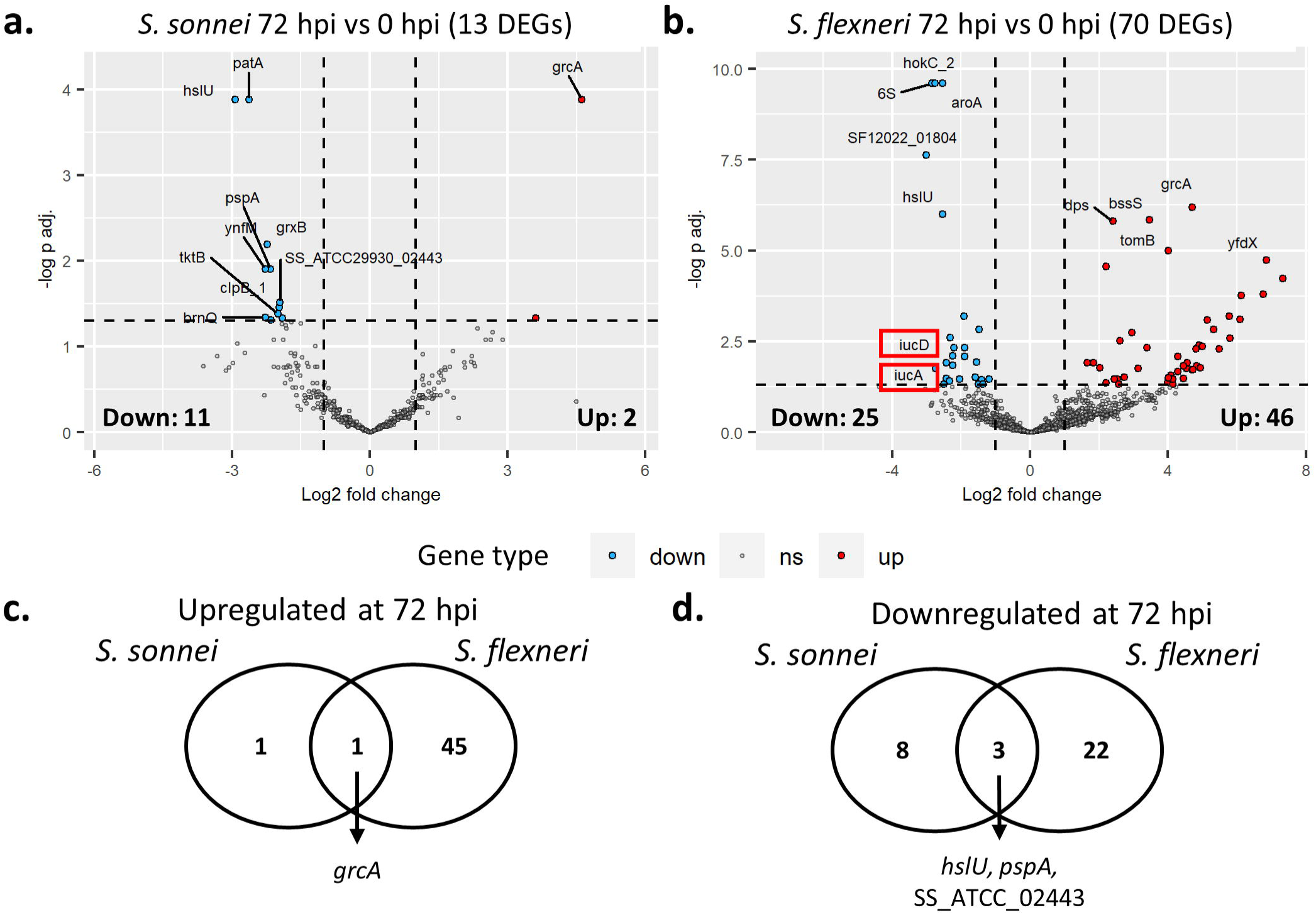
Comparison between *S. sonnei* 29930 and *S. flexneri* 12022 responses at 72 hpi and 0 hpi of *C. elegans* infection (72 hpi vs 0 hpi). **a.** Volcano plot (-log10(p-adjusted value) versus log2 fold change) of the differentially expressed genes (DEGs) in *S. sonnei* at 72 hpi compared to 0 hpi. **b.** Volcano plot of the DEGs in *S. flexneri* at 72 hpi compared to 0 hpi. The virulence genes *iucA* and *iucD* are indicated in red boxes. Significantly downregulated genes are shown in blue, and significantly upregulated genes are shown in red. Non-significant genes are shown in grey. **c.** The downregulated DEGs in both *S. sonnei* and *S. flexneri* at 72 hpi. d. The upregulated DEGs in both *S. sonnei* and *S. flexneri* at 72 hpi.

As no other virulence genes are differentially regulated at 72 hpi, the other DEGs may provide more information on *S. sonnei* and *S. flexneri* response during *C. elegans* infection. No enriched pathways are identified, indicating that the responses of these two strains during infection involved only individual genes, and not any specific pathways. Both species commonly upregulate 1 gene (*grcA*) and downregulate 3 genes (*hslU, pspA,* SS_ATCC_02443) during infection (Figure 5c, 5d). The glycyl radical cofactor A (*grcA*) is a cofactor that restores the activity of oxygen-damaged pyruvate formate-lyase (PflB), an enzyme important for anaerobic glycolysis [73]. The GrcA protein expression is enhanced by acidic pH and low levels of oxygen, conditions which are common within the gut [74]. However, this contradicts another study that reported the downregulation of *grcA* by *S. sonnei* during zebrafish infection [23]. The downregulation of *hslU, pspA,* and SS_ATCC_02443 (hypothetical protein) genes in both species suggests their non-crucial role in *S. sonnei* and *S. flexneri* during *C. elegans* infection. The *hslU* gene encodes for an ATP-dependent protease that plays a role in unfolding proteins for degradation [75]. At higher temperatures, this protease will degrade proteins and increases the rate of ATP hydrolysis [76]. The phage shock protein A PspA prevents secretin pores misinsertion in *E. coli* and has been implicated in *Salmonella* virulence [77]. The hypothetical protein was identified as *hok,* the toxin gene of type I *hok*-*sok* toxin-antitoxin (TA) system that forms pores in the bacterial membrane and leads to cell death [78]. Among these 3 genes, *pspA* was reported to be upregulated by *S. sonnei* during zebrafish infection [23] and *S. flexneri* during HeLa cells infection [24], which contradicts this study. The discrepancy of *grcA* and *pspA* expression may be due to the different animal models and conditions during infection, as this study looked into the response of *Shigella* in *C. elegans* after 72 hours at 25 °C, while Torraca *et al.* investigated *S. sonnei-*infected zebrafish larvae for 24 hours at 28.5 °C [23]. The regulation of these genes may be specific to the animal model and infection conditions.

Other than *grcA, S. sonnei* upregulates *malP,* which is not seen in *S. flexneri.* MalP, a maltodextrin phosphorylase, plays a key role in maltose/maltodextrin and glycogen metabolism [79, 80]. While *malP* mutants do not have a negative impact on pathogenic *E. coli* colonization, the ability to metabolize maltose provides an advantage for *S. sonnei* during intestinal colonization [81]. In contrast, *S. flexneri* upregulates a broader range of genes, including the biofilm regulator *bssS,* DNA-binding protein *dps,* Hha toxicity modulator *tomB,* and a protein with unknown function *yfdX* (Figure 5b). In particular, a number of genes related to acid resistance or tolerance are upregulated in *S. flexneri* (*gadABCE, hdeAB, frc, ydeP*). *Shigella* have multiple mechanisms to survive in acidic conditions in the host gut, including the amino acid decarboxylase/antiporter system that depends on glutamate (GadAB/GadC) [82]. The upregulation of these genes in *S. flexneri* and not *S. sonnei* suggests that *S. flexneri* induces the expression of these genes at 72 hpi, while *S. sonnei* may have constantly upregulated these genes at the beginning of infection. This is observed during the direct comparison between *S. sonnei* and *S. flexneri* at 0 hpi, where the genes *gadC* and *hdeA* are significantly upregulated in *S. sonnei* compared to *S. flexneri* (Supplementary Table 6). The low number of DEGs in *S. sonnei* than *S. flexneri* also suggests that *S. flexneri* regulates its response towards infection after infection began, while *S. sonnei* may be constantly regulating the necessary genes to promote its survival regardless of the infection timepoint.

### *S. sonnei* upregulates *fepA* and other non-virulence genes during *C. elegans* infection compared to *S. flexneri*

Next, the differential gene expression analysis was performed between *S. sonnei* 29930 and *S. flexneri* 12022 at the same time of infection (*S. sonnei* vs *S. flexneri*). Only the 3366 genes that are present in both strains are considered in this analysis to allow the direct comparison between these two strains. There are 45 DEGs identified at 0 hpi between *S. sonnei* and *S. flexneri*, where 39 and 6 genes are upregulated and downregulated respectively (Figure 6a; Supplementary Table 6). These genes were differentially expressed even before infection was established, thus serving as the baseline differences between these two strains. At 72 hpi, 214 genes are differentially expressed between *S. sonnei* and *S. flexneri*, where 129 genes are upregulated, and 85 genes are downregulated (Figure 6b; Supplementary Table 7).

**Figure 6:**
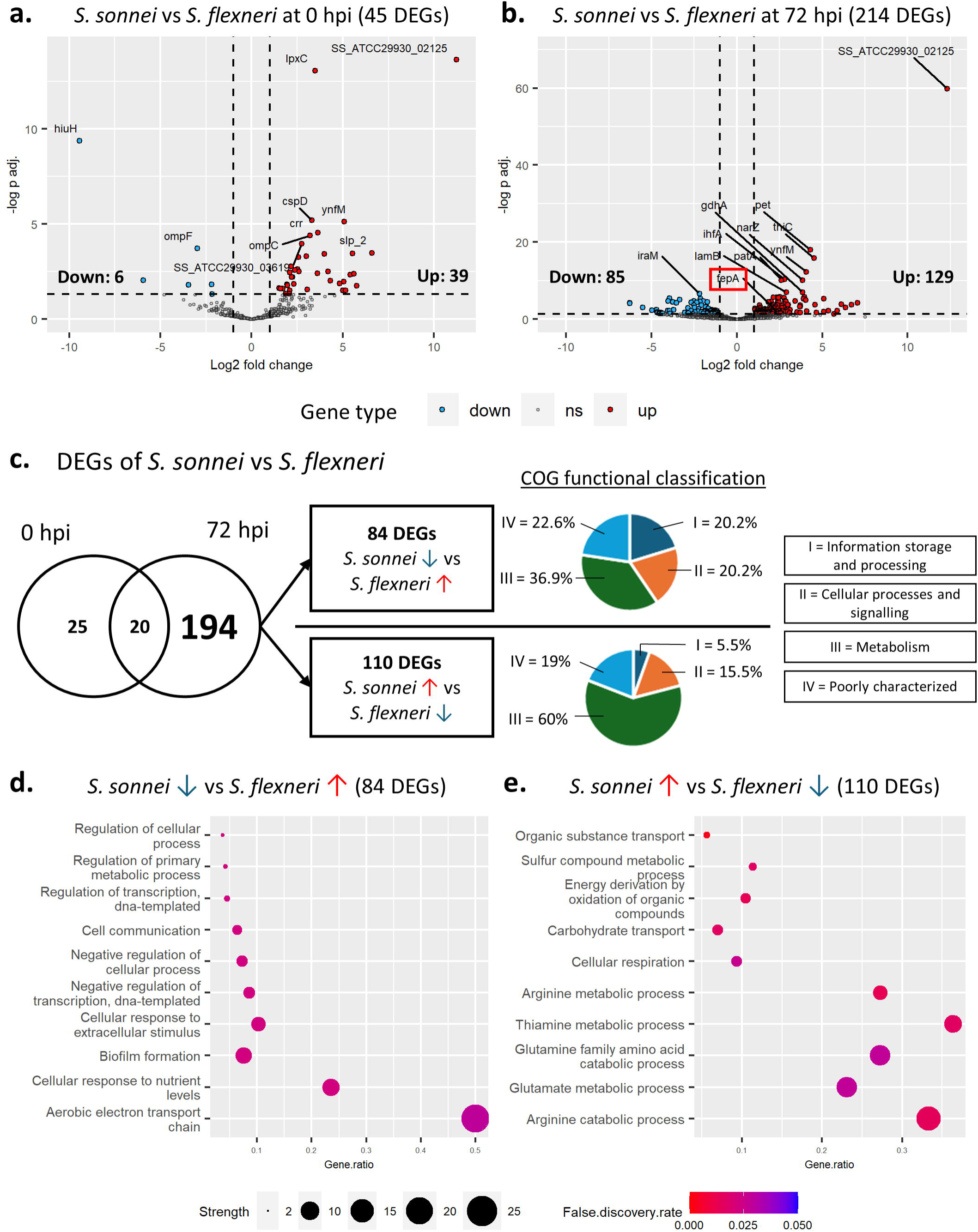
Comparison between *S. sonnei* and *S. flexneri* responses at the same time of *C. elegans* infection (*S. sonnei* vs *S. flexneri*). **a.** Volcano plot (-log10(p-adjusted value) versus log2 fold change) of the differentially expressed genes (DEGs) at 0 hpi between *S. sonnei* and *S. flexneri*. **b.** Volcano plot of the DEGs at 72 hpi between *S. sonnei* and *S. flexneri*. The virulence gene *fepA* is indicated with a red box. Significantly downregulated genes are shown in blue, and significantly upregulated genes are shown in red. Non-significant genes are shown in grey. **c.** The DEGs of *S. sonnei* and *S. flexneri* at the two different timepoints, and the COG classification of the 194 genes related to infection only. **d.** The enriched pathways in the 84 downregulated DEGs in *S. sonnei* vs *S. flexneri* comparison. **e.** The enriched pathways in the 110 upregulated DEGs in *S. sonnei* vs *S. flexneri* comparison. Red up arrow (↑) indicates upregulation, and blue down arrow (↓) indicates downregulation.

Among the 24 common virulence genes within these two strains (Figure 4a), only one virulence gene was significantly differentially regulated (Figure 6b, highlighted in red box). The ferric enterobactin outer membrane transporter gene *fepA* was significantly upregulated by *S. sonnei* 29930 compared to *S. flexneri* 12022 at 72 hpi of *C. elegans* infection. The *fep* genes play a role in ferric enterobactin uptake during iron utilization, and *fepA* specifically encodes a TonB-dependent outer-membrane receptor that actively transports ferric enterobactin into the periplasm of the bacteria [55, 83]. Similar to this study, the *fepA* gene is upregulated in *S. sonnei* after 24 hpi of zebrafish larvae infection [23]. The *fepA* gene, like the other enterobactin genes, is conserved in all 46 (100%) *S. sonnei* strains analyzed in the pangenome analysis, while 91 (82%) *S. flexneri* strains have this gene (Figure 3b). As discussed previously, enterobactin is a part of *S. sonnei’s* repertoire in virulence that is not utilized in *S. flexneri* [59]. This study revealed that among the genes in enterobactin synthesis and transport, *fepA* plays an important role in *S. sonnei* during *C. elegans* infection.

Among the significantly upregulated DEGs in *S. sonnei* at both 0 hpi and 72 hpi is the *lpxC* gene (Figure 6a, Supplementary Table 6 and 7). This gene encodes for the UDP-3-O-acyl-N-acetylglucosamine deacetylase gene involved in the biosynthesis of lipid A within the LPS layer [84]. Additionally, the hypothetical gene SS_ATCC29930_02125 (identified as a general stress protein) is also consistently upregulated at both time points by *S. sonnei*. The amino acid sequence of this stress protein is 100% identical to *E. coli* YmfD protein, which is upregulated by *E. coli* in response to multiple stressors (heat, antibiotics, and oxidative stress) [85]. While its specific function is unknown, the upregulation of this gene suggests that *S. sonnei* is responding to the *C. elegans* infection as a form of stress, which is not directly seen in *S. flexneri*.

### *S. sonnei* upregulates metabolism genes while *S. flexneri* upregulates non-metabolism genes during *C. elegans* infection

There are 20 genes that are differentially regulated between *S. sonnei* and *S. flexneri* at both timepoints. To further investigate the genes that are only differentially expressed during 72 hpi of *C. elegans,* these 20 genes are removed from further analysis. The remaining 194 genes were split into 84 DEGs that are downregulated in *S. sonnei,* and 110 DEGs that are upregulated in *S. sonnei* (Figure 6c). Their COG categories were determined using eggNOG v5.0 (Figure 6c). A functional enrichment analysis using Gene Ontology (GO) terms using ShinyGo v0.80 (with the Gene Ontology Biological Processes category) was performed on these genes (Figures 6d and 6e).

COG categorization analysis demonstrated a distinct distribution of the gene functions between genes upregulated by *S. sonnei* and *S. flexneri* (Figure 6c). The COG categories are placed into 4 groups: information storage and processing (I = J, K, L), cellular processes and signaling (II = D, M, N, O, T, U, V), metabolism (III = C, E, F, G, H, I P, Q), and those poorly or not characterized (IV) genes. The distribution of gene functions in the 84 DEGs upregulated by *S. flexneri* are evenly distributed between these 4 groups. However, within the 110 DEGs upregulated by *S. sonnei*, 60% of them are metabolism genes. This suggests that during *C. elegans* infection, *S. sonnei* upregulates more metabolism genes and pathways than *S. flexneri*.

The functional pathways enriched within the 84 DEGs upregulated by *S. flexneri* are mostly cellular processes and transcription regulation (Figure 6d). In particular, 6 genes are involved in all of these regulation pathways, which are *farR, yefM, ebgR, yhcK, cpxP, and zur*. Other pathways include cell communication (*phoP, phoQ, appA, iraM, yedW, rcsB, dinD, yhjX, yjiY*), and response to extracellular stimulus (*appA, iraM, dinD, yhjX, yjiY*). Compared to *S. sonnei, S. flexneri* may be upregulating these regulatory genes to fine-tune its response towards infection instead of regulating the metabolic genes directly. Some genes also reflect the acid tolerance response of *S. flexneri* at 72 hpi, such as the *phoPQ* genes. These genes are involved in a two-component system that allows tolerance towards acidic pH within the host environment [86], and thus virulence [87]. Additionally, *S. flexneri* upregulates genes involved in biofilm formation (*ybaJ, yefM, ycfR, yjgK*), and aerobic electron transport chain (*appB, appC*), indicating their importance in *S. flexneri* pathogenesis. The expression of the biofilm regulator YbaJ, the outer membrane protein YcfR, the antitoxin YefM, and the protein YjgK have been shown to promote biofilm formation in *E. coli* [88–90]. In *S. flexneri,* biofilm is induced by bile-salt within the gut, and it leads to increased bacterial cell aggregation during infection [91]. While the biofilm of *S. sonnei* is also induced by bile-salts, the importance of *S. sonnei* biofilm during infection is not determined. Next, the aerobic electron transport chain genes, *appB* and *appC genes* encode for a cytochrome bd-II oxidase. These genes are advantageous for *E. coli* during intestinal inflammation in murine model [92]. The upregulation of these biofilm formation and aerobic electron transport chain genes in *S. flexneri* and not *S. sonnei* suggests that they play a larger role in *S. flexneri* response during infection.

In contrast, among the 110 genes upregulated by *S. sonnei* during *C. elegans* infection, the organic substance transport, multiple metabolic processes (including sulfur compound, arginine, thiamine, glutamine, and glutamate), and cellular respiration are enriched (Figure 6e). The metabolic processes include the arginine catabolic process (*astA, astB, speA*), thiamine metabolic process (*thiH, thiG, thiE, thiC*), and sulfur compound metabolic process (*acs, cysD, cysN, cysQ, metE, sbp, tauD, ynjE*). This suggests that *S. sonnei* may prioritize their metabolism and growth during infection by scavenging or synthesizing important nutrients, or trigger metabolic changes in the host to maintain their infection [93]. Sulfur is needed in amino acids (e.g. cysteine and methionine) and other cofactors such as iron-sulfur clusters, thiamine, and coenzyme A [94]. Thus, the acquisition and assimilation of the limited sulfur within the host is key for the continued survival of the bacteria. Other than sulfur, thiamine is also required for pathogenesis and bacterial growth during infection [95, 96].

Additionally, thiamine and arginine provide acid tolerance or resistance [97, 98]. These upregulated metabolic processes suggest an adaptive response towards low pH conditions during gut infection by *S. sonnei* but not *S. flexneri* [99],. Lastly, *S. sonnei* upregulates genes involved in organic substance transport and carbohydrate transport, including mannose transport (*manX, manY, manZ*), dipeptide transport (*dppA*, *dppD, dppF*), and galactose transport (*mglB, mglC*). The increase in substrate transport indicates the increase in metabolism within *S. sonnei,* supporting the idea that *S. sonnei* mainly upregulates metabolism during infection compared to *S. flexneri* (Figure 6c).

### The pangenome, virulence gene profiles, and transcriptomic response during *C. elegans* infection of *S. sonnei* is distinct from *S. flexneri*

In this study, pangenome and transcriptomic analysis of *S. sonnei* and *S. flexneri* were performed to elucidate the causes of the increased *S. sonnei* prevalence. Pangenome analysis revealed that *S. sonnei* has a more conserved genome than *S. flexneri*, which is more diverse (Figure 2c and 2d). Overall, *S. sonnei* has more virulence genes than *S. flexneri* (Figure 3d). 19% of *S. sonnei* virulence genes are core genes (present in 100% of the strains), but *S. flexneri* has no core virulence genes (Figure 3e). The highly conserved virulence genes in *S. sonnei* strains include G4C operon (*yccC, yccY, yccZ, ymcA, ymcB, ymcC,* and *ymcD*), MAM gene, T6SS (*fha, hcp1, hcp2, rhs, tssA, tssB, tssF, tssG, tssJ, tssL, tssM,* and *tssI*), enterobactin synthesis and transport genes (*entA, entB, entC, entD, entE, entF, fepA, fepB, fepC, fepD, fepG*), aerobactin synthesis and transport genes (*iucA, iucB, iucC, iucD, iutA*), and ferrous iron transport (*sitA, sitB,* sitD). Next, the *C. elegans* infection model was studied using the two representative strains *S. sonnei* ATCC 29930 and *S. flexneri* ATCC 12022, where *S. sonnei* is more virulent than *S. flexneri* even with similar bacterial load (Figure 4c and 4d). *S. sonnei* differentially regulates less DEGs than *S. flexneri.* This suggests that while *S. flexneri* respond to infection by differentially regulating a higher number of genes, *S. sonnei* is more equipped for infection even before infection is established, and thus do not need to differentially regulate a lot of genes. Other than that, the analysis on the 194 infection-specific DEGs between *S. sonnei* and *S. flexneri* revealed that *S. sonnei* upregulates 110 DEGs that includes *fepA,* a virulence gene involved in ferric enterobactin uptake, and other genes involved in sulfur, thiamine, and arginine metabolism. In contrast, *S. flexneri* upregulates 84 DEGs that are involved in aerobic electron transport chain, and the regulation of transcription, biofilm formation, cell communication, and response to extracellular stimuli.

Overall, *S. sonnei* is shown to prioritize the uptake and utilization of essential but limited resources during infection (i.e. iron, sulfur, thiamine, arginine), and these genes are likely to be constantly expressed. The presence of chromosomal core virulence genes in *S. sonnei* also suggests that this species have similar response and strategies regardless of the strain, however this needs further validation. In contrast, *S. flexneri* prioritizes the ability to regulate the cell’s response only when it is needed, such as during 72 hpi. As *S. flexneri* is more diverse, other strains of *S. flexneri* may have different strategies during infection, such as the use of other virulence genes absent in *S. flexneri* ATCC 12022. The differences between *S. sonnei* and *S. flexneri* observed in this analysis are depicted in Figure 7.

**Figure 7:**
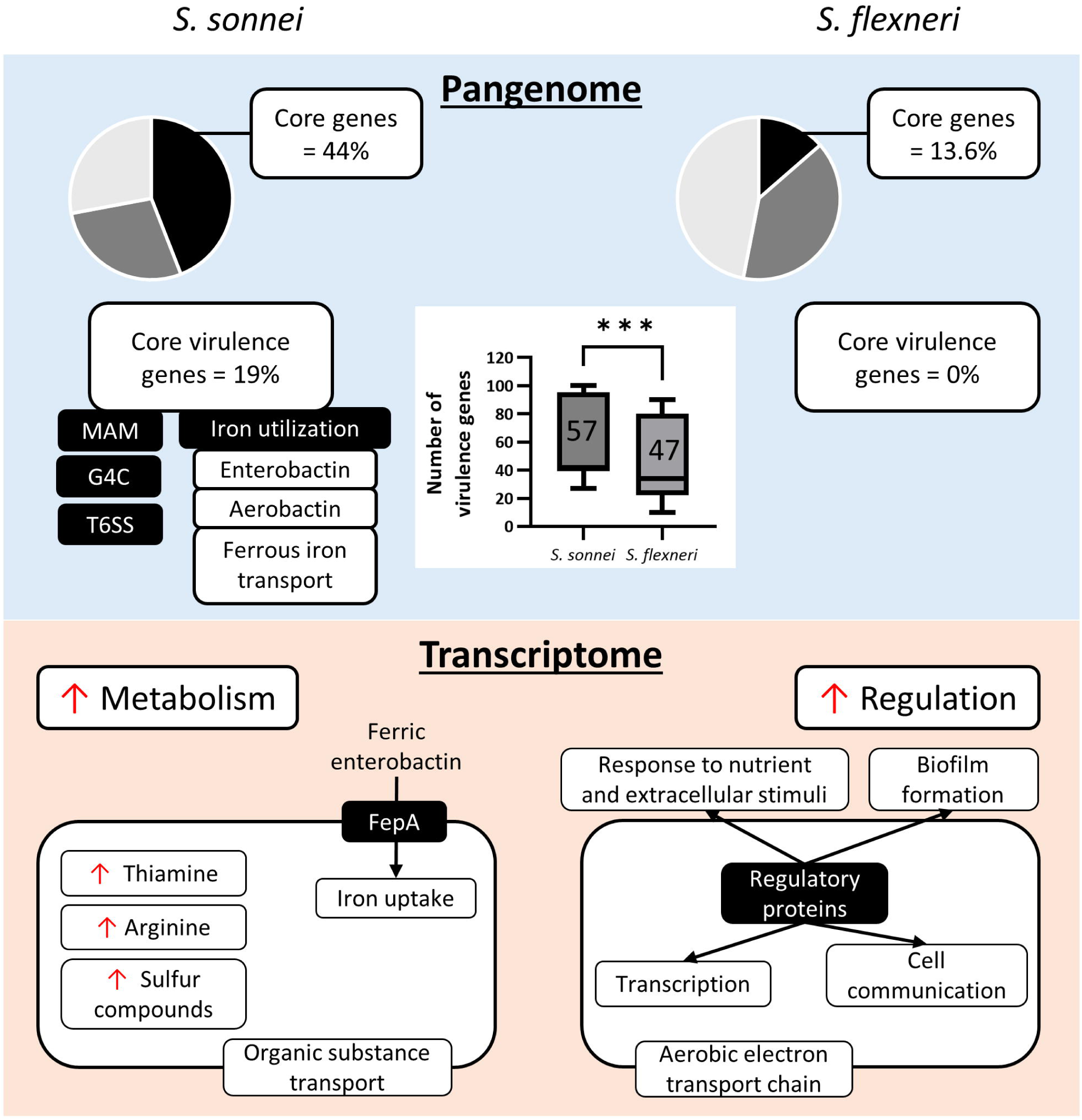
Schematic diagram of the species-specific differences between *S. sonnei* and *S. flexneri* revealed from pangenome and transcriptomic analysis.

There are some limitations in this study. The virulence genes analyzed in this study may not be an exhaustive list, as novel virulence genes or genes that are newly associated with *S. sonnei* or *S. flexneri* virulence may be discovered. Next, a low number of virulence genes are detected as DEGs. This is likely due to their low gene expression. Due to the nature of dual-RNA sequencing, the current experimental setup only allows the detection of highly expressed genes in bacteria, and thus the genes with low expression may not be detected in this study. Additionally, the infection was performed using *C. elegans,* which is maintained at 25 °C. *Shigella,* as a human pathogen, thrives at 37 °C (human body temperature) instead. Subsequent validation of our findings should be performed in another model that is closer to human physiological conditions.

## Conclusion

In conclusion, this study compares the pangenome and transcriptomic response between *S. sonnei* and *S. flexneri* and revealed multiple differences between these two species. Firstly, the higher number of core genes in *S. sonnei* than *S. flexneri* indicates the conserved genome of *S. sonnei* and the diversity of *S. flexneri* strains. The number of virulence genes is significantly higher in *S. sonnei* than in *S. flexneri. S. sonnei* has core virulence genes (19%), but not *S. flexneri.* These core virulence genes include the G4C, MAM, T6SS, and multiple iron utilization genes (encoding for enterobactin, aerobactin, and ferrous iron transport). Transcriptomic analysis of two representative *Shigella* (*S. sonnei* ATCC 29930 and *S. flexneri* ATCC 12022) revealed that these two *Shigella* species employ different strategies during *C. elegans* infection at 72 hpi. Firstly, *S. sonnei* differentially expressed less genes during infection than *S. flexneri*. It is proposed that while *S. flexneri* regulates its response towards *C. elegans* after infection begins, *S. sonnei* instead constantly express the genes necessary for infection even before infection begins. Next, direct comparison between the responses of these two species revealed that while *S. sonnei* upregulates metabolism-related genes and pathways (sulfur, thiamine, arginine), *S. flexneri* instead upregulates genes involved in regulation of transcription, biofilm, and other cellular process. The virulence genes contributed minimally to virulence in these two strains as most of them are not differentially regulated during infection. However, *S. sonnei* significantly upregulates *fepA,* the outer membrane protein responsible for ferric enterobactin uptake during infection as compared to *S. flexneri.* Overall, the presence of core virulence genes, and the upregulation of metabolism and the virulence gene *fepA* in *S. sonnei* may be one of the factors leading to *S. sonnei* slowly overtaking *S. flexneri* as the cause of shigellosis. Further studies are needed to confirm and validate their distinct roles in *S. sonnei* and *S. flexneri*.

## Abbreviations

ANOVA: Analysis of variance
ATCC: American Type Culture Collection
CDS: coding sequences
CFU: Colony forming units
COG: Clusters of Orthologous Genes
DEG: Differentially expressed genes
FC: Fold change
FDR: False discovery rate
G4C: Group 4 capsules
GF: Gene Family
GO: Gene Ontology
hpi: hours post infection
LB: Luria-Bertani
LMIC: Low- and middle-income countries
LPS: Lipopolysaccharide
MAM: Multivalent adhesion molecule
ML: M9 buffer containing 25 mM levamisole
MLG: M9 buffer containing 25 mM levamisole and 1mg/ml of gentamycin
NGM: Nematode growth medium
OD: Optical density
PCA: Principal component analysis
ShET1: *Shigella* enterotoxin 1
SPATE: Serine protease autotransporters of Enterobacteriaceae
T3SS: Type III secretion system
T6SS: Type VI secretion system
TA: Toxin-antitoxin
VFDB: Virulence Factor Database
WRAIR: Walter Reed Army Institute of Research

## Author contributions

WBC: experimental design, acquisition and interpretation of data, writing – drafting, review & editing, WWY: Interpretation of data, writing - review & editing, THS: supervision, conceptualization, writing – review & editing

## Data availability

Trimmed paired-end data of the dual RNA sequencing (*Caenorhabditis elegans* - *Shigella sonnei* and *Caenorhabditis elegans* - *Shigella flexneri* at 72 hours post-infection) are available from the Sequence Read Archive (SRA) under BioSample SAMN40006912 (*S. sonnei*) and SAMN40006913 (*S. flexneri)*, in the BioProject PRJNA1078575. Similar data for 0 hours post-infection (control) is available under BioSample SAMN42050357 (*S. sonnei*) and SAMN42050358 (*S. flexneri)* in the BioProject PRJNA1128419. Other datasets used and analyzed during the current study are available from the corresponding author upon reasonable request.

