## Supplementary figures and images for "Pangenome and Transcriptomic Analysis Revealed Distinct Virulence Genes Profiles and Infection Responses in *Shigella Sonnei* and *Shigella Flexneri*"

### Supplementary Fig 1

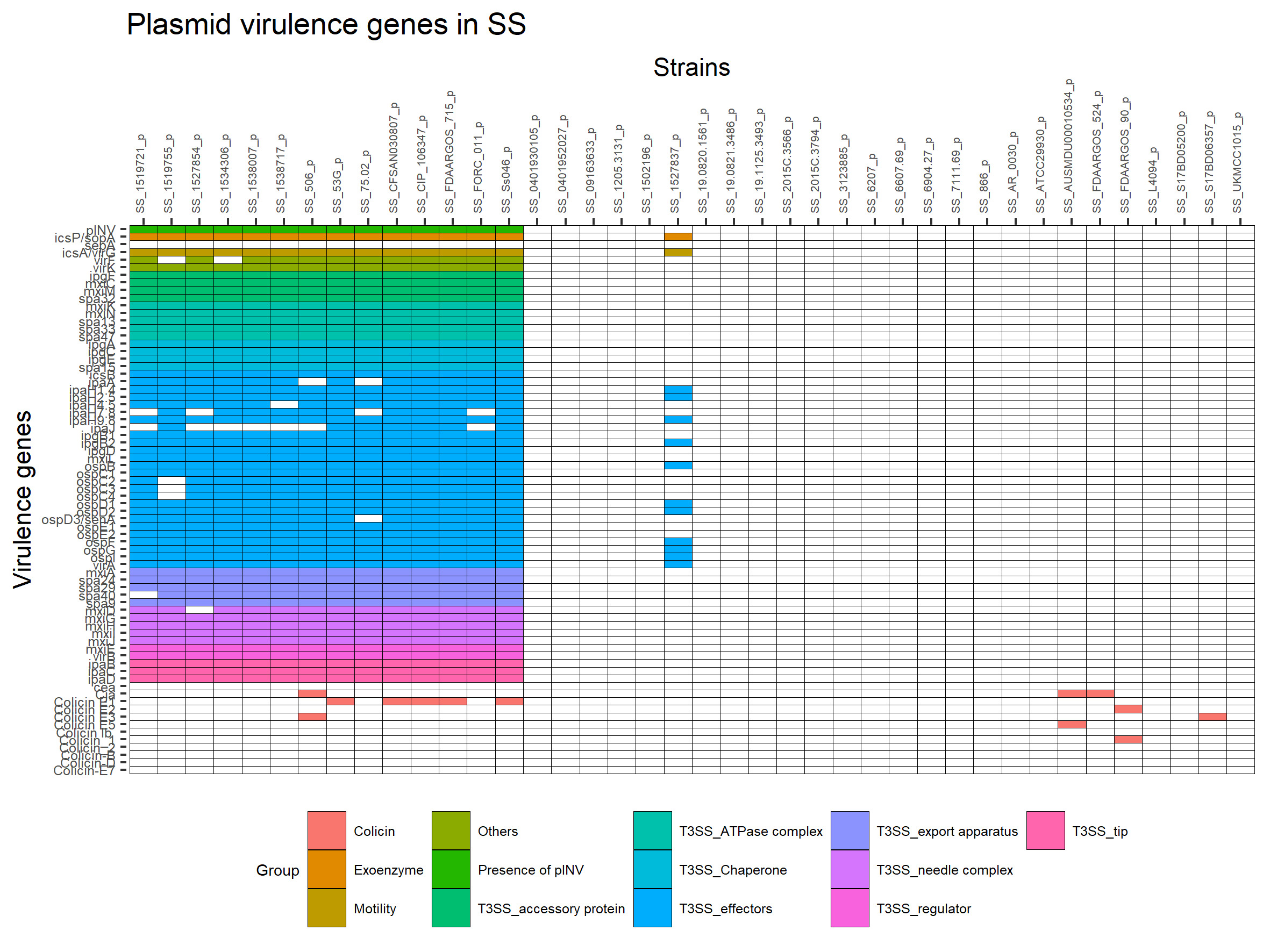

### Supplementary Fig 2

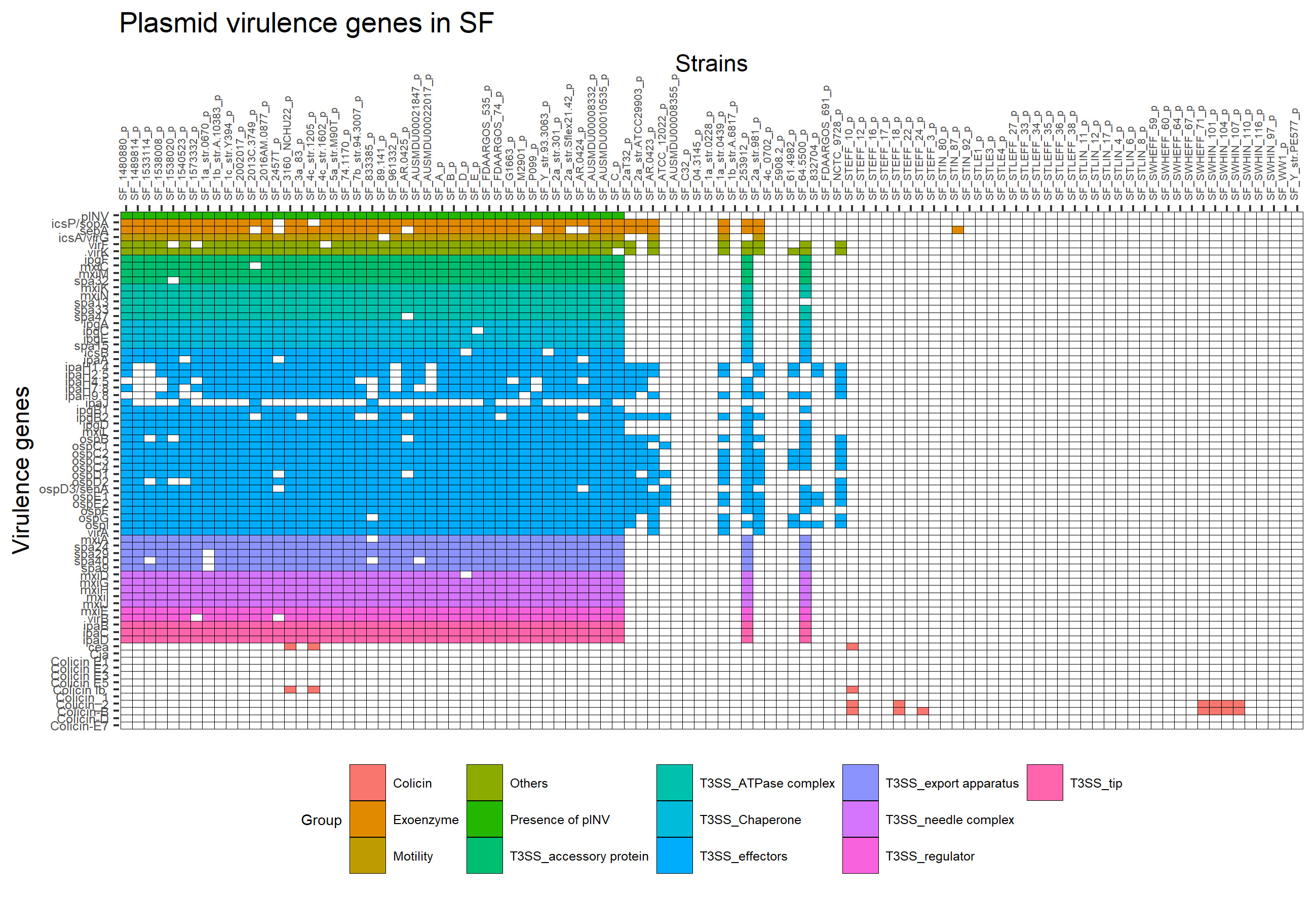

### Supplementary Fig 3

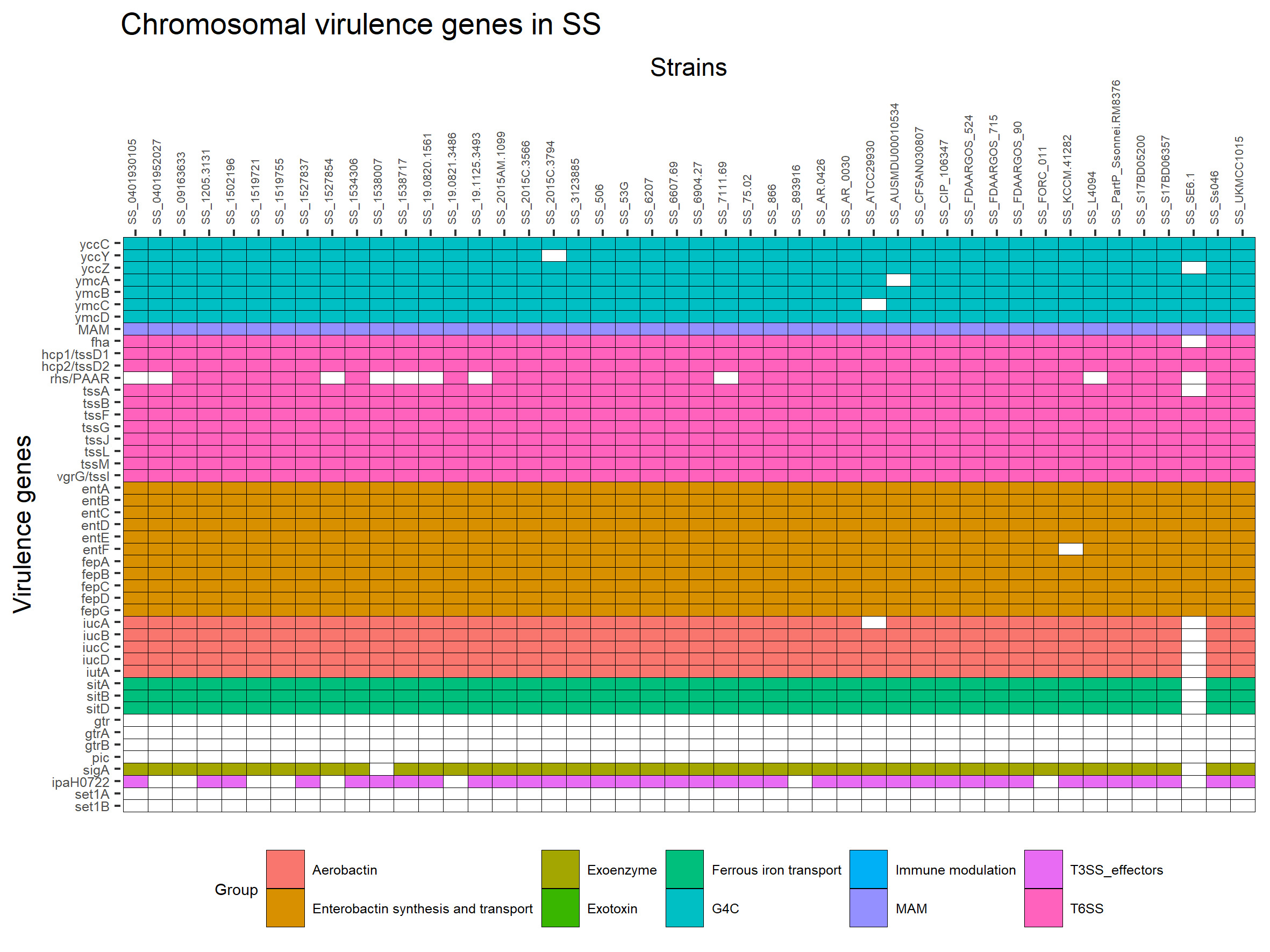

### Supplementary Fig 4

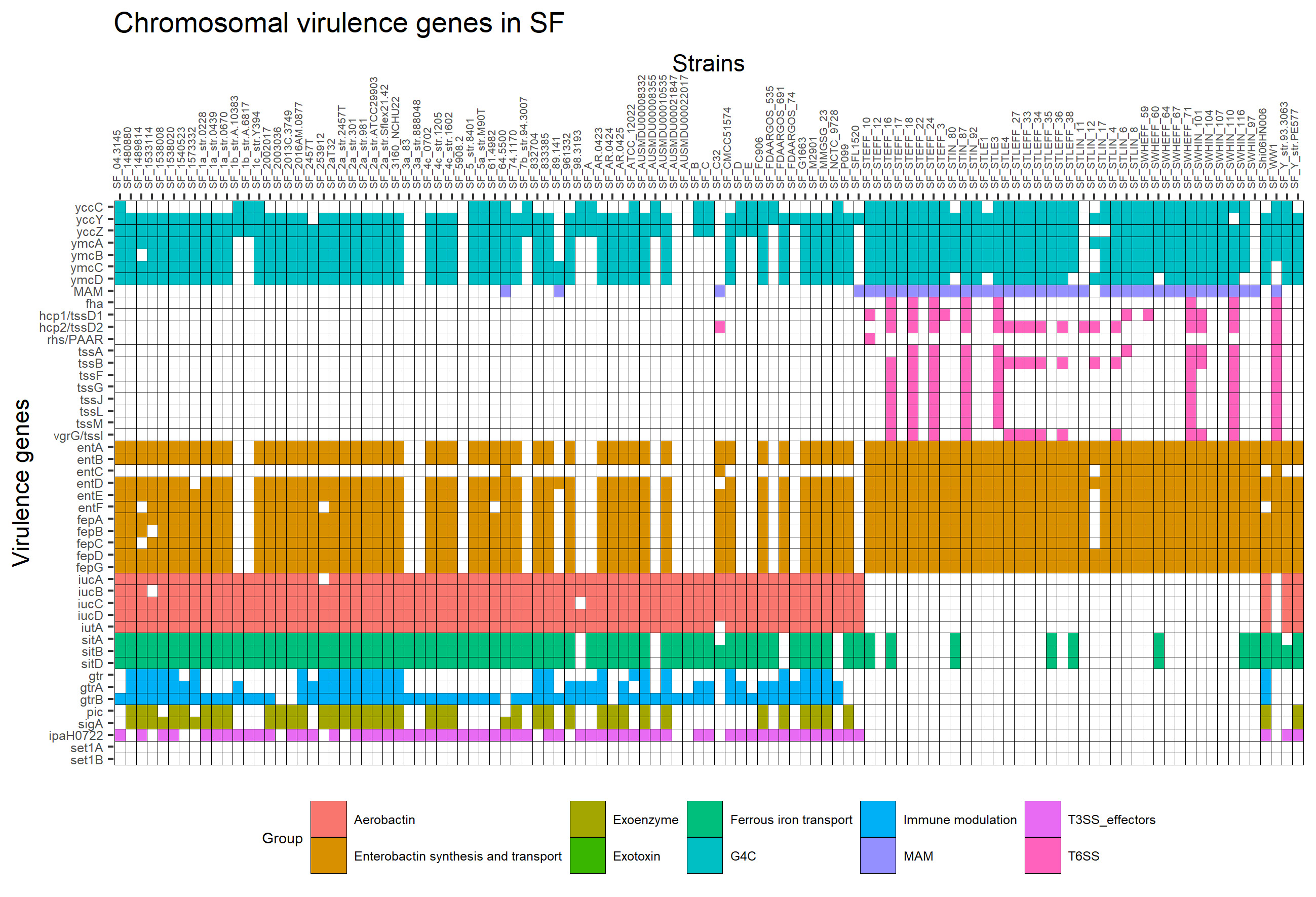

### Supplementary Fig 5

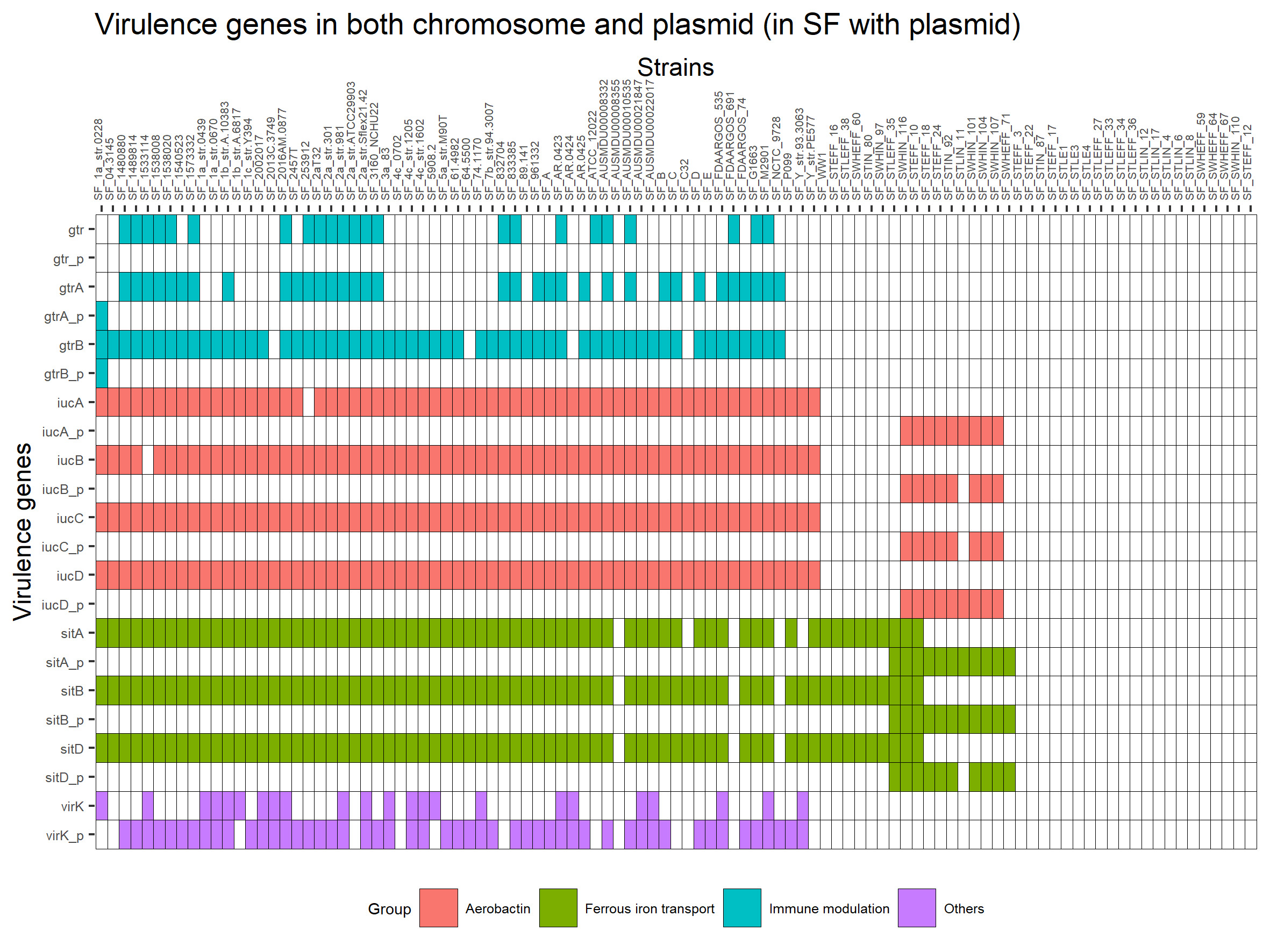

### Supplementary Fig 6

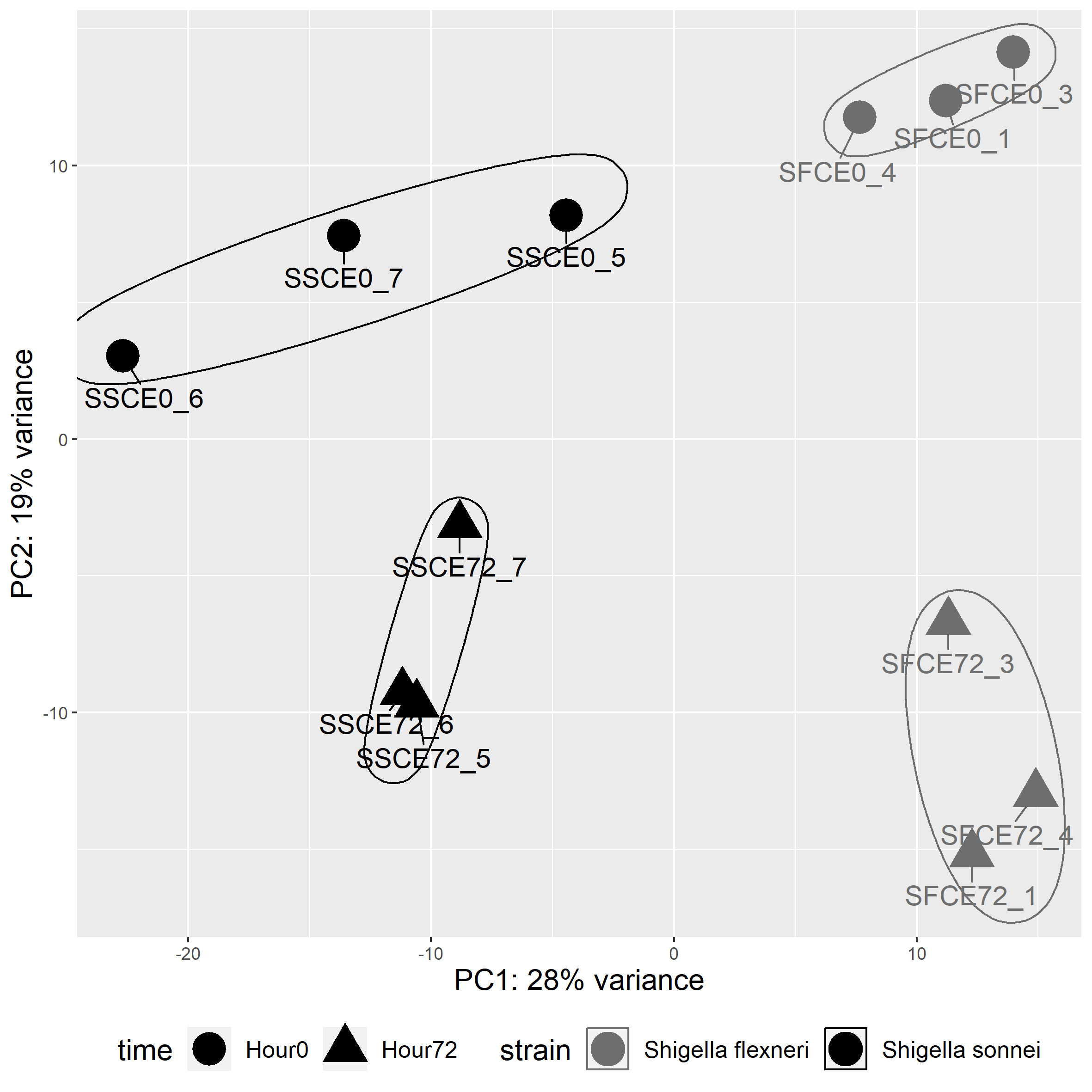
